# The cyanotoxin 2,4-DAB enhances mortality and causes behavioral and molecular dysfunctions associated with neurodegeneration in larval zebrafish

**DOI:** 10.1101/2021.10.13.464292

**Authors:** Rubia M. Martin, Michael S. Bereman, Kurt C. Marsden

## Abstract

Exposure to cyanotoxins has been linked to neurodegenerative diseases, including amyotrophic lateral sclerosis, Alzheimer’s, and Parkinson’s disease. While the cyanotoxin β-methylamino-L-alanine (BMAA) has received much attention, cyanobacteria produce many cyanotoxic compounds, several of which have been detected in nature alongside BMAA including 2,4-diaminobutyric acid (2,4-DAB), and N-(2-aminoethyl)glycine (AEG). Thus, the question of whether DAB and AEG also cause neurotoxic effects *in vivo* is of great interest, as is the question of whether they interact to enhance toxicity. Here, we evaluate the toxic and neurotoxic effects of these cyanotoxins alone or in combination by measuring zebrafish larval viability and behavior after exposure. 2,4-DAB was the most potent cyanotoxin as it decreased larval viability by approximately 50% at 6 days post fertilization, while BMAA and AEG decreased viability by just 16% and 8%, respectively. Although we only observed minor neurotoxic effects on spontaneous locomotion, BMAA and AEG enhanced acoustic startle sensitivity, and they interacted in an additive manner to exert their effects. 2,4-DAB, however, only modulated the startle kinematics, an indication of motor dysfunction. To investigate the mechanisms of 2,4-DAB’s effects, we analyzed the protein profile of larval zebrafish exposed to 500μM 2,4-DAB at two time points and identified molecular signatures consistent with neurodegeneration, including disruption of metabolic pathways and downregulation of the ALS-associated genes SOD1 and UBQLN4. Together, our data demonstrate that BMAA and its isomers AEG and 2,4-DAB cause neurotoxic effects *in vivo*, with 2,4-DAB as the most potent of the three in the zebrafish model.

## 1. Introduction

Exposure to BMAA (β-N-methylamino-L-alanine), a non-canonical amino acid, has been extensively associated with the onset of sporadic neurodegenerative diseases, including amyotrophic lateral sclerosis (ALS) (Banack and Cox 2003), Parkinson’s (Nunes-Costa et al. 2020) and Alzheimer’s diseases (Silva et al. 2020). BMAA is naturally released into the water by a diverse taxa of cyanobacteria (Cox et al. 2005), and it persists in the environment not only during and after a cyanobacteria bloom collapse (Réveillon et al. 2015) but also in terrestrial ecosystems (Metcalf et al. 2012). BMAA can also bioaccumulate and be transferred to higher trophic levels (Li et al. 2019). *In vitro* studies have shown that BMAA causes excitotoxicity in primary neuronal and glial cell cultures (Chiu et al. 2012; 2013), and it is also known to induce differential expression of genes and proteins associated with cellular mitochondrial dysfunction and protein aggregation (Beri et al. 2017; Chiu et al. 2015). These results suggest that BMAA is a potent epidemiological link to sporadic neuropathology. Not only is BMAA found ubiquitously in the environment, but it can also be found along with other highly prevalent cyanotoxins, including BMAA derivatives like 2,4 diaminobutyric acid (2,4-DAB) and aminoethyl glycine (AEG) (Chatziefthimiou et al. 2018; Metcalf et al. 2008; Réveillon et al. 2015). However, the neurotoxicity of these isomers is largely unknown, and because a natural exposure is likely to involve a combination of BMAA, its isomers, and other co-occurring cyanotoxins, it is particularly important to investigate the neurotoxic effects of exposure to cyanotoxic mixtures.

A preliminary binary mixture study showed synergistic neurotoxic effects of low concentrations of BMAA (10-100μM) and methylmercury (3μM) in primary cortical cell cultures (Rush et al. 2012). We have also previously demonstrated how BMAA can interact with isomers (i.e., 2,4-DAB and AEG) in an *in vitro* system to enhance caspase activity and induce neurodegenerative processes in motor neuron-like cells (NSC-34) (Martin et al. 2019). A recent *in vitro* study revealed that the isomer AEG is a more potent neurotoxin than BMAA and that the toxicity of AEG is mediated by the induction of free radicals as well as the activation of metabotropic glutamate receptors (i.e., mGluR5) (Schneider et al. 2020). Like BMAA, 2,4-DAB activates N-methyl-D-aspartate (NMDA) receptors (Spasic et al. 2018), a mode of action this is also a primary mechanism of BMAA-induced neurotoxicity (Lobner 2009). These *in vitro* studies confirm the neurotoxicity of 2,4-DAB and AEG and highlight the importance of examining their potential combined effects with BMAA *in vivo*.

Here we aimed to identify which individual cyanotoxic isomer and/or which combination of BMAA, AEG, and 2,4-DAB elicits the greatest toxicity in larval zebrafish, using an axial simplex mixture design (Cornell 2011). Here, we evaluated the larval zebrafish viability after exposure to 10 different ratios of all three cyanotoxic components. We also evaluated behavioral indicators of neurotoxicity with a high-throughput behavior testing platform and found that there was little interaction between the toxins, and that 2,4-DAB is a more potent toxin than either BMAA or AEG *in vivo*. To then identify early molecular changes that trigger toxicity of 2,4-DAB we performed discovery proteomics at two developmental timepoints after exposure. Our results show that processes associated with neurodegeneration are impacted, including Ca^2+^ signaling, the unfolded protein response, and endoplasmic reticulum stress. Together, our data highlight the importance of studying mixtures and reveal 2,4-DAB to be a more potent neurotoxin than BMAA and AEG in the zebrafish model.

## 2. Materials and Methods

### 2.1 Chemicals

Synthetic BMAA, AEG and 2,4-DAB standards were obtained from Sigma Aldrich (St. Louis, MO). Water, acetonitrile, methanol, acetic acid, and formic acid were all Optima LC–MS grade solvents purchased from Fisher Scientific (Tewksbury, MA, USA). Stock solutions of BMAA, AEG, and 2,4-DAB at 10 mg.mL-1 were used for all mixture’s dilutions. All dilutions were prepared in HPLC grade water.

### 2.2 Mixture Design

Experiments were designed to address *in vivo* toxicity after exposure to BMAA, AEG, and 2,4-DAB alone or in combination, in which both variation in experiment replicates and reproducibility were assessed. To detect interactive effects in a three-component mixture we used simplex axial design (Cornell 2011). Three compounds were studied at 7 different mixture combinations. The simplex axial design generally includes a test solution named center-point in which all factors are studied at their equimolar concentrations and the linear average value between their maximum (1) and minimum (−1) levels, allowing for an accurate estimation of pure error, lack-of-fit (Cornell 2011). The mixture design was generated using Design Expert® software where each run was controlled to a total concentration of 500μM. The 500μM concentration were chosen based on the lowest observable adverse effect level (LOAEL) found in our previous zebrafish dose response studies (Martin et al. 2020).

### 2.3 Zebrafish husbandry and exposures

All animal use and procedures were approved by the North Carolina State University IACUC. Zebrafish (*Danio rerio*) embryos from multiple crosses of wild type tupfel longfin (TLF) strain adults were collected and placed into Petri dishes containing E3 medium, and unfertilized eggs were removed as described previously. Embryos from all clutches were mixed and randomly sorted into 24 well plates (8-10 animals per well) containing 1 mL of E3 per well.

At 6 hours post fertilization (hpf), embryos were treated with vehicle, individual cyanotoxins, their binary mixtures, and their three-component mixtures (**Figure 1**). All treatments were performed in triplicate and were repeated in each of 2 separate experiments. Embryos were incubated at 29°C on a 14h:10h light-dark cycle, and 100% of the media was exchanged for fresh solutions daily. Embryos/ larvae were exposed to treatments until 6 days post fertilization (6 dpf).

**Figure 1.**
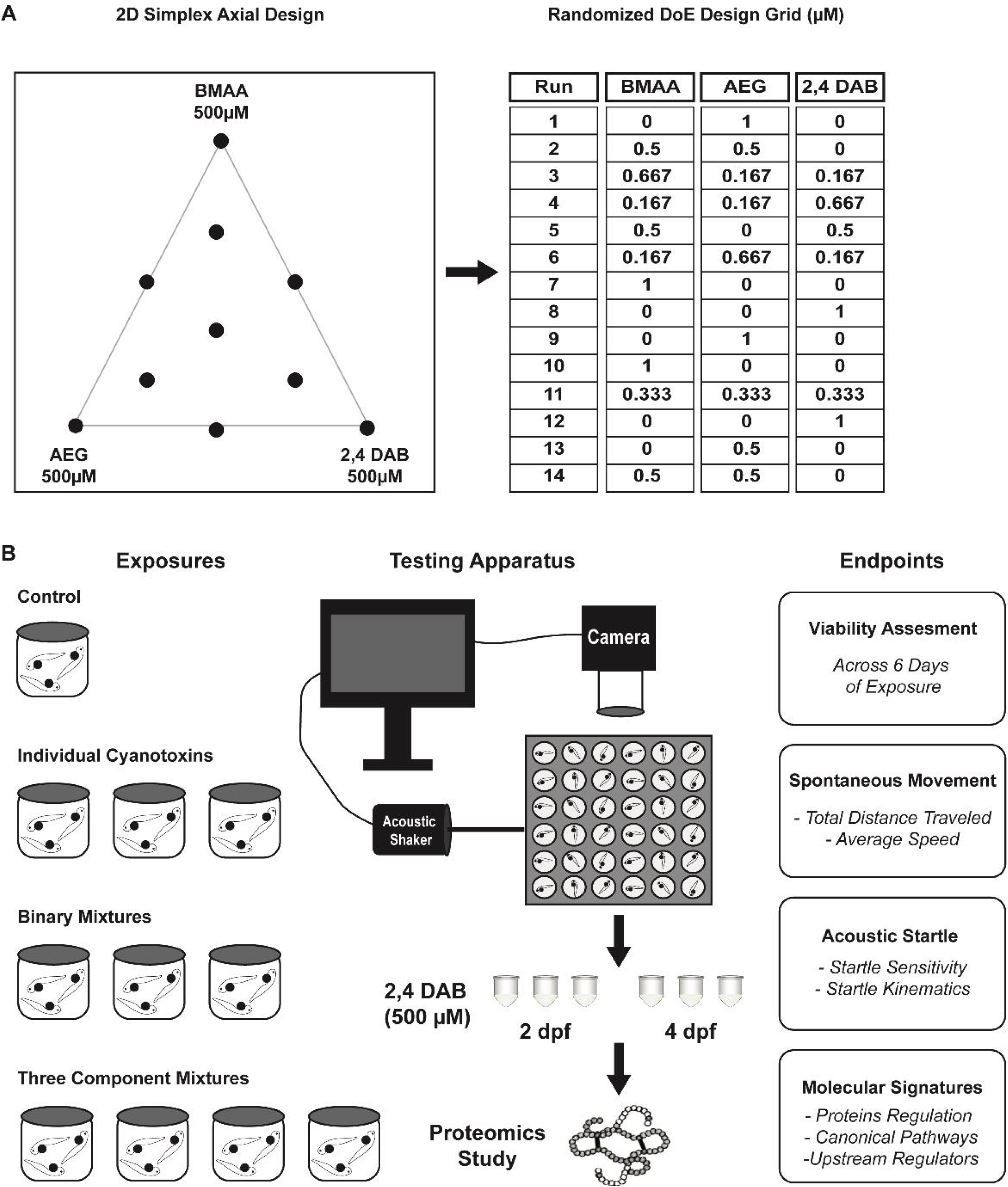
Experimental design. 2D simplex axial design and randomized Design of Experiments(DoE) design grid representing the composition for each solution. (B) Exposure plan for larval zebrafish from 6 hours post fertilization (hpf) to 6 days post fertilization (dpf) followed by the high throughput behavior testing, proteomics analysis, and endpoints measurements for these studies.

### 2.4 Zebrafish Viability and Behavior Assessment

Mortality due to treatment toxicity was assessed daily and dead fish were counted and removed. During daily media changes, each fish was also assessed for a set of developmental phenotypes (pericardial edema, otolith malformations, pigmentation defects, small eyes, small heads, body axis defects such as curved or bent tails, and uninflated swim bladders), and any fish exhibiting these phenotypes was removed. Significant differences in viability data were analyzed via Kaplan-Meir survival analysis with Mantel-Cox Log-rank test using Prism 8.

### 2.5 Behavior assays and analysis

Screened 6 dpf larvae were adapted to the testing lighting and temperature conditions for 30 minutes prior to testing. Behavior testing was done as previously described (Burgess and Granato 2007a; Marsden et al. 2018). Briefly, larvae were transferred to individual 9 mm round wells on a 36-well laser-cut acrylic testing grid. Larvae acclimated for 5 min and then spontaneous locomotor activity was recorded for 18.5 min at 640 x 640 px resolution at 50 frames per sec (fps) using a Photron mini UX-50 high-speed camera. The same set of larvae were then presented with a total of 60 acoustic stimuli, 10 at each of 6 intensities (13.6, 25.7, 29.2, 35.5, 39.6, and 53.6 dB), with a 20 s interstimulus interval (ISI). Startle responses were recorded at 1000 fps. Stimuli were delivered by an acoustic-vibrational shaker to which the testing grid was directly mounted. All stimuli were calibrated with a PCB Piezotronics accelerometer (#355B04) and signal conditioner (#482A21), and voltage outputs were converted to dB using the formula dB = 20 log V. Analysis of recorded behaviors was done using FLOTE software as described previously (Marsden et al. 2018). Startle response C-bends were automatically identified using defined kinematic parameters. A startle sensitivity index was calculated for individual larvae by calculating the area under the curve of startle frequency versus stimulus intensity using Prism 8 software (GraphPad). Statistical analyses were performed using JMP pro 14 from SAS institute, Cary, NC. Data were analyzed for effects between the groups (comparison of means), using Tukey-Kramer HSD, Alpha 0.05. Bar graphs were generated using Prism 8.

### 2.6 Global Proteomics Study

#### 2.6.1 The cyanotoxin 2,4-DAB exposure

Zebrafish embryos from at least ten different fish pairs per batch were collected and randomized immediately after fertilization and transferred in groups of 10 into 24-well plates in E3 media. At 6 hpf, zebrafish embryos were treated with vehicle or 2,4-DAB at 500μM in E3. Embryos were incubated at 29°C on a 14h:10h light-dark cycle, and 100% of the media was exchanged for fresh treated solutions daily. Embryos/ larvae were exposed to 2,4-DAB until 2 or 4 dpf. Ten larvae were pooled for each of 3 biological replicates for each condition and then flash frozen in liquid nitrogen and stored at −80°C.

#### 2.6.2 Sample preparation and LC MS/MS

Details of sample preparation, protein extraction and digestion via filter aided sample preparation (FASP) can be found in Supplemental Methods. Details regarding the LC-MS/MS data collection are also provided in the Supplemental Methods. Raw data files obtained in this experiment have been made available on the Chorus LC-MS data repository and can be assessed under the project ID#1738.

#### 2.6.3 Proteomics Data Analysis

Label free quantification (LFQ) was performed in MaxQuant (version.1.5.60), which uses the Andromeda algorithm (Tyanova et al. 2016). Both dynamic (i.e., methionine oxidation and N-terminal acetylation) and fixed modifications (i.e., cysteine carbamido-methylation) were used for the database search along with a maximum of two missed cleavages. The minimum peptide length was set to 7 amino acids and the false discovery rate (FDR) for peptide and protein identification was set to 0.01. All other search parameters were left as default values. Data were searched against the *Danio rerio* Swiss Prot protein database (# protein sequences = 56 281, accessed 01/25/2021). Comparison of LFQ intensities across the whole set of measurements was investigated using Perseus software (version 1.5.1.6), where calculation of statistical significance was determined using two-way Student’s t-test and FPR (p value ≤0.05).

#### 2.6.4 Pathway Analysis

Ingenuity Pathway Analysis (IPA) software was used to identify the function, specific processes, and enriched pathways of the differentially expressed proteins using the “Core Analysis” function. Only significantly differentially expressed proteins (p value ≤ 0.05) were submitted to IPA. We used an empirical background protein database to evaluate the significance of pathway enrichment. The database was created by using all the proteins that were detected in our samples (Khatri and Drăghici 2005).

## 3. Results

An overview of the experiments in this study is illustrated in **Figure 1**. An optimal mixture design was created using Design of Experiments (DoE) to establish the concentrations of BMAA, AEG, and 2,4-DAB to be used in the exposures. The 2D axial design generates a symmetrical triangle plot for a three-variable case, displaying 7 mixtures of different ratios along with each individual cyanotoxin. An additional 4 replicate tests were applied for a total of 14 runs to evaluate reproducibility and the lack of fit for the derived model (**Figure 1A**). In brief, zebrafish larvae were exposed to BMAA, AEG, and 2,4-DAB as well as their binary/three-component mixture combinations from 6 hpf to 6 dpf. Viability of the larvae was assessed, and neurotoxicity was evaluated via two behavioral assays: spontaneous locomotion and acoustic startle response. To investigate perturbed molecular pathways associated with 2,4-DAB toxicity, the 2,4-DAB-exposed larvae were collected and subjected to discovery proteomics at two developmental stages: 2 dpf and at 4 dpf (**Figure 1B**).

### 3.1 Viability Assessment: 2,4-DAB is more toxic than BMAA and AEG in vivo

To determine if exposure to cyanotoxins could induce organism mortality, we aimed to evaluate the viability of zebrafish larvae during exposure to BMAA, AEG, and 2,4-DAB alone or in combination. We exposed zebrafish larvae from 6 hpf to 6 dpf to cyanotoxic solutions as illustrated in **Figure 1**. We did not observe any significant overt developmental phenotypes in any of the exposed larval zebrafish groups. We performed Kaplan-Meir survival analysis with a Mantel-Cox Log-rank test to determine if BMAA, AEG, and 2,4-DAB significantly alter survival over time. While there was a trend toward a slight decrease in viability, AEG did not significantly decrease larval survival when compared to the control group (P=0.0531). However, larval survival after exposure to either BMAA or 2,4-DAB was significantly reduced by 16% (P=0.0008) and 50% (P<0.0001), respectfully (**Figure 2A**). Surprisingly, 2,4-DAB was the most potent cyanotoxin, and its EC50 for induced death is 500μM. The survival model standard errors, coefficients, and probability of survival values are included in **Table 1**. These data indicate that 2,4-DAB is highly toxic to zebrafish larvae at moderately low concentrations.

**Table 1.** Fitted Survival Probability and Coefficient Intervals.

| Treatment | Std Error | Lower 95% | Upper 95% | Prob Survival |
| --- | --- | --- | --- | --- |
| Control | 0.01224 | 0.00173 | 0.08226 | 0.98768 |
| BMAA | 0.04095 | 0.09618 | 0.25833 | 0.83856 |
| AEG | 0.02914 | 0.03372 | 0.15543 | 0.92581 |
| 2,4-DAB | 0.04682 | 0.41143 | 0.59294 | 0.49774 |

**Figure 2.**
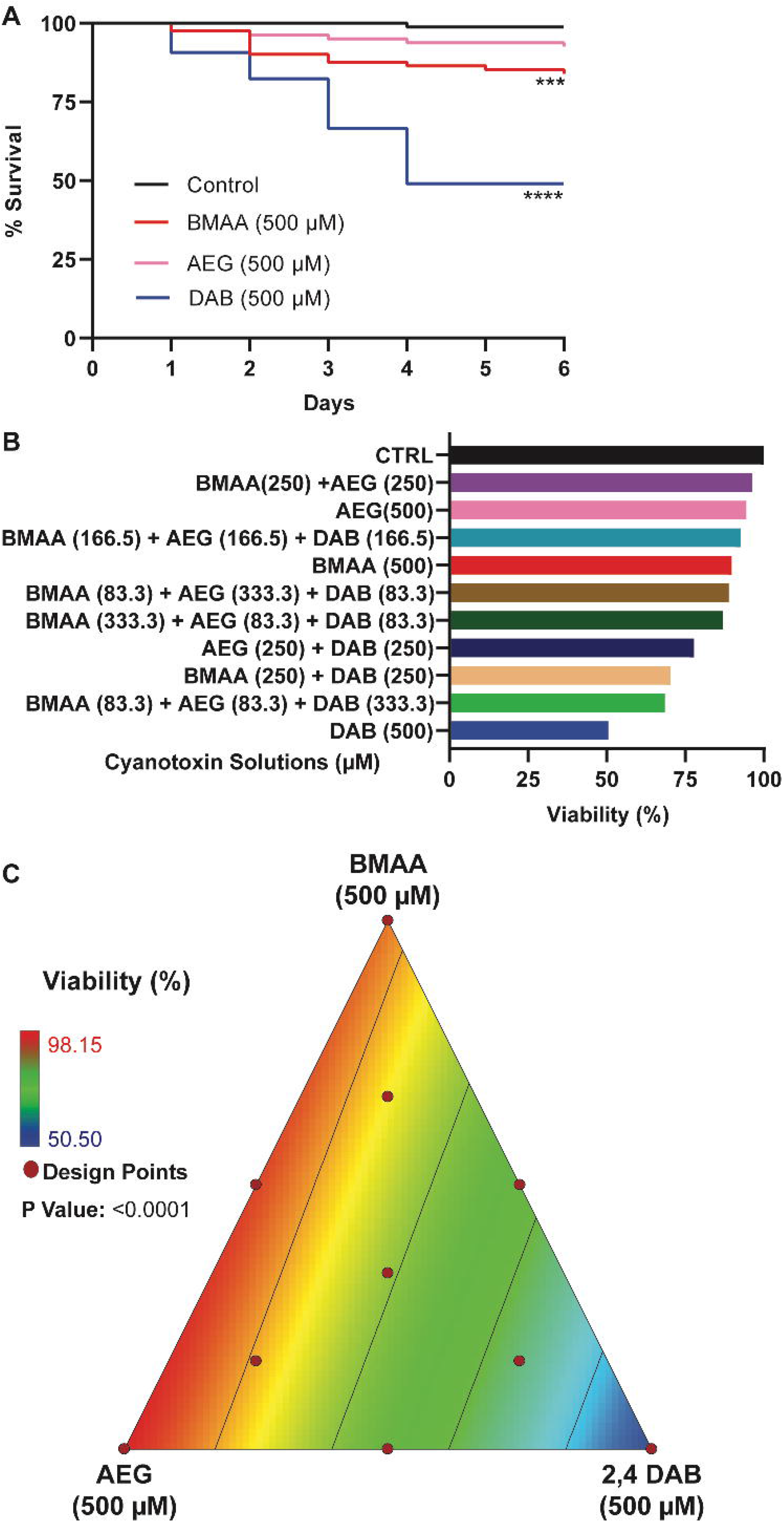
Viability Assessment. (A) Kaplan-Meier cumulative survival plots comparing percent survival between four treatment groups: Control (n=108), BMAA (n=162), AEG (n=162), and 2,4-DAB (n=108) (P<0.001, Mantel-Cox log rank test). (B) Bar graph representing percent viability across all treatment groups. (C) 2D contour plot representing zebrafish percent viability in response to each mixture of BMAA, AEG and 2,4-DAB (Linear regression model, P<0.0001).

When we assessed the survival of zebrafish larvae exposed to the binary and three-component mixtures of BMAA, AEG, and 2,4-DAB, we observed a trend towards reduced viability (i.e., ranging from 11.2% to 31.5%) for the larval group exposed to the cyanotoxic mixtures containing increasing concentrations of 2,4-DAB (**Figure 2B**). Thus, 2,4-DAB alone drives the reduced viability of zebrafish larvae exposed to cyanotoxic mixtures, and when combining it with other cyanotoxin, toxicity is not enhanced. Then, we tested if there existed a significant relationship between 2,4-DAB and decreased viability using the simplex axial design. **Figure 2C**illustrates the viability across all mixture conditions, and it indicates a plot of viscosity as a function of all three-mixture components, BMAA, AEG and 2,4-DAB. The grid lines representing percent viability decease in value towards the 2,4-DAB vertex. At 500μM, 2,4-DAB exhibited the lowest viability (**Figure 2C**). The analysis of variance indicates that a linear model is statistically significant (P<0.0001) (**Table 2**).

**Table 2.** Analysis of Variance (ANOVA): Viability. ANOVA results, showing the significance a linear model (DF – degrees of freedom).

|  | Sum of Squares | DF | Mean Square | F Value | P Value |
| --- | --- | --- | --- | --- | --- |
| **Model** | 3205.63 | 2 | 1602.82 | 68.01 | < 0.0001 |
| **Linear Mixture** | 3205.63 | 2 | 1602.82 | 68.01 | < 0.0001 |
| **Residual** | 259.24 | 11 | 23.57 |  |  |
| **Lack of Fit** | 202.66 | 7 | 28.95 | 2.05 | 0.2549 |
| **Pure Error** | 56.58 | 4 | 14.15 |  |  |

Thus, we conclude that these three cyanotoxins do not interact to decrease survival *in vivo* and that 2,4-DAB is more toxic than BMAA and AEG, meaning the viability response tracks linearly with 2,4-DAB alone. The response is modeled by the below linear model in which *Y_i_* represents the predicted response, β ο is the intercept coefficient, and β_*i*_ is the coefficient of the linear regression.

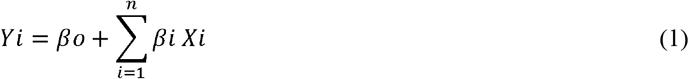

### 3.2 Mixture of cyanotoxins modulate spontaneous behavior in larval zebrafish

To determine if exposure to BMAA, AEG, and 2,4-DAB alone or in combination causes neurotoxicity in larval zebrafish, we first examined spontaneous locomotion across all 10 treatment groups (**Figure 3**). The surviving zebrafish larvae at 6 dpf were adapted to the testing conditions, and then their spontaneous movements were recorded for 18.5 min using a high-throughput behavior platform and automated FLOTE tracking software (Martin et al. 2020). We detected no significant differences in total distance travelled for larvae treated with BMAA, AEG, or 2,4-DAB compared to their respective vehicle controls (**Figure 3A**). Thus, exposure to BMAA, AEG, or 2,4-DAB alone does not grossly impact motor function. Although the average speed of the BMAA and 2,4-DAB-exposed groups was not altered when compared to their respective controls, the average speed of the larvae treated with AEG was significantly reduced (**Figure 3A**). The binary mixtures containing AEG also decreased spontaneous movement, with AEG plus BMAA reducing speed and AEG plus 2,4-DAB decreasing both distance travelled and speed (**Figure 3B**). We also observed that exposure to the equimolar three-component mixture significantly decreased both distance travelled and average speed (**Figure 3C**). DOE models were unable to fit these data at a statistically significant level, precluding any determination of whether there were interactions between toxins in this assay. These data indicate that although individual concentrations of BMAA and AEG do not substantially reduce survival of larval zebrafish, combined they can induce toxicity at a behavioral level.

**Figure 3.**
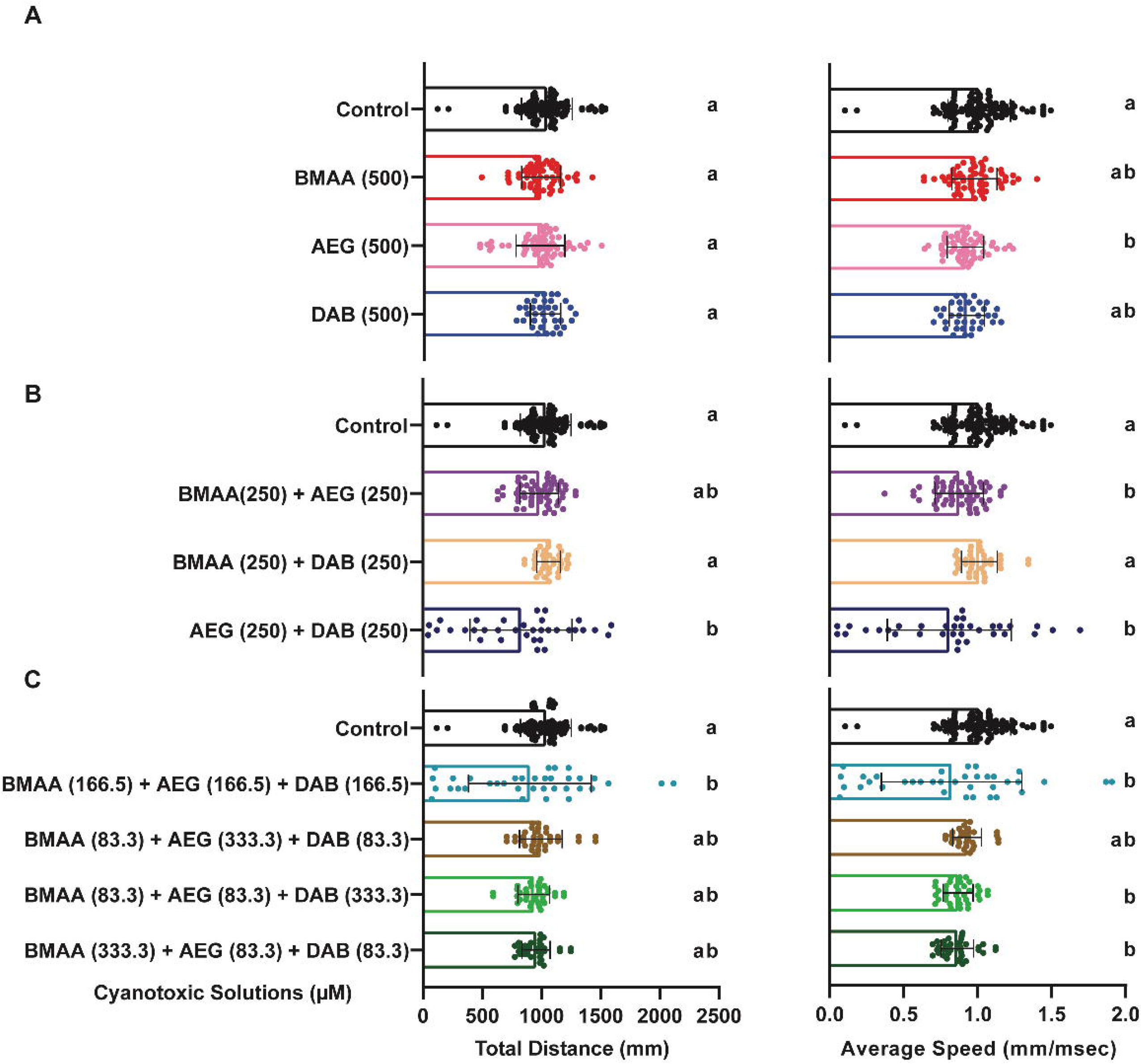
Spontaneous Locomotion Response. (A - C) Bar graphs represent the total distance travelled during the 18.5 min spontaneous movement assay for each larva and average speed across the same assay for BMAA, AEG, and 2,4-DAB alone, their binary mixtures and three-component mixtures, respectively. Levels not connected by the same letter are significantly different (Tukey-Kramer HSD, Alpha 0.05).

### 3.3 The cyanotoxin 2,4-DAB does not alter acoustic startle sensitivity, but BMAA and AEG do

To determine if exposure to BMAA, AEG, and 2,4-DAB alone or in combination affects specific sensorimotor behaviors in larval zebrafish, we measured acoustic startle frequency in response to a range of stimulus intensities in the same 10 groups described above. The frequency of Mauthner cell-dependent short latency C-bends (SLCs) are displayed in **Figure 4 A-C**. We previously found that BMAA increases SLC frequency, suggesting increased excitability of the startle circuit (Martin et al. 2020). Here, we found that like BMAA, AEG also increased SLC frequency, but 2,4-DAB did not (**Figure 4A**). We also observed that the binary mixture of BMAA and AEG caused a similar increase in SLC frequency compared to BMAA and AEG alone (**Figure 4B**). No other binary or three component mixture induced statistically significant effects compare to controls. We also analyzed the frequency of long latency C-bends (LLCs), which are normal responses typically elicited by weaker stimuli and that are independent of the Mauthner cell and instead rely on the activation of prepontine neurons (Marquart et al. 2019). Neither BMAA nor its isomers significantly altered LLC response frequency (**Supplemental Figure 1**). These data indicate that BMAA and AEG specifically enhance the sensitivity of the SLC circuit. We next modeled the SLC experiment by fitting the same linear equation described above. **Figure 4D** shows the grid lines representing the SLC response across all 10 tested mixtures. SLC sensitivity increased towards the pure doses of BMAA (500μM) and AEG (500μM) as well as their binary mixture (BMAA at 500μM plus AEG at 500μM). The analysis of variance indicates that the linear model is statistically significant (P<0.0001) (**Table 3**). These data reveal that BMAA and AEG alone can augment the SLC response and that they interact, likely additively, at lower concentrations to significantly enhance SLC responses in larval zebrafish. These data also reveal that 2,4-DAB alone or in combination with BMAA and/or AEG does not affect the initiation of the SLC response.

**Table 3.** Analysis of Variance (ANOVA): Short Latency C Startle. ANOVA results, showing the significance a linear model (DF – degrees of freedom).

|  | Sum of Squares | DF | Mean Square | F Value | P Value |
| --- | --- | --- | --- | --- | --- |
| **Model** | 2.148E+005 | 2 | 1.074E+005 | 5.20 | 0.0257 |
| **Linear Mixture** | *2.148E+005* | 2 | *1.074E+005* | *5.20* | *0.0257* |
| **Residual** | 2.271E+005 | 11 | 20641.63 |  |  |
| **Lack of Fit** | *1.253E+005* | 7 | *17906.09* | *0.70* | *0.6790* |
| **Pure Error** | *1.017E+005* | 4 | *25428.83* |  |  |

**Figure 4.**
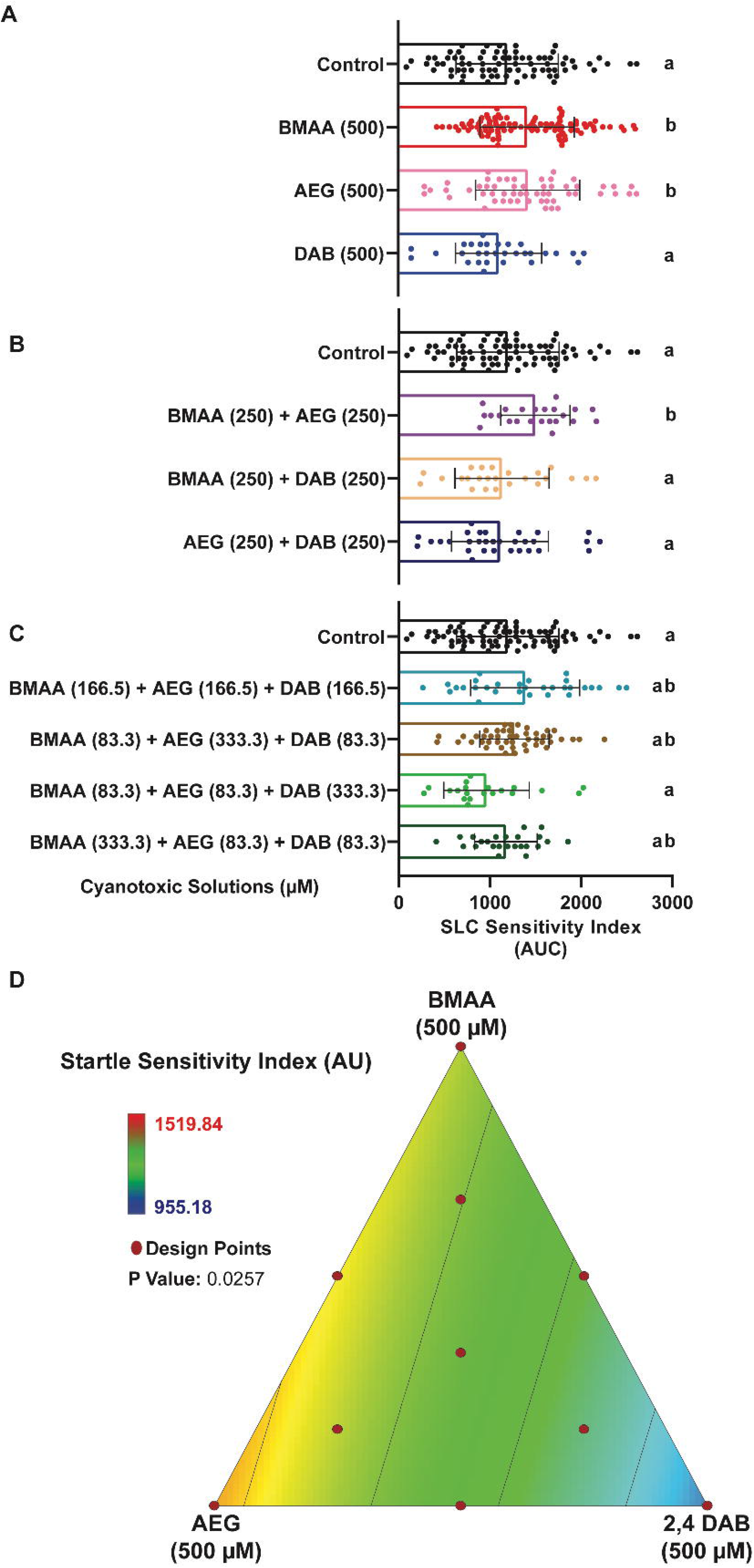
BMAA and AEG augment the short latency c startle (SLC) response. (A-C) Bar graphs display the distribution of the short latency C-bend (SLC) sensitivity indices for each tested larva after exposure to individual cyanotoxins, binary mixtures and three component mixtures, respectively. SLC sensitivity index is determined for each fish by calculating the area under the curve of SLC frequency vs. stimulus intensity (n = 108 siblings; mean ± SEM). Levels not connected by the same letter are significantly different (Tukey-Kramer HSD, Alpha 0.05). (D) 2D contour plot for the SLC sensitivity at different points in the design space, including each individual cyanotoxin (BMAA, AEG and 2,4-DAB) and their seven different mixture ratios (Linear regression model, P=0.0257).

### 3.4 The cyanotoxin 2,4-DAB modulate acoustic startle kinematics in larval zebrafish

To assess whether cyanotoxin mixtures cause more subtle defects in motor control, we examined the kinematic performance of SLC responses in the exposed larvae, as these rapid movements require coordinated activation of large sets of motor neurons and muscle. Using FLOTE tracking software, we looked at a set of kinematic parameters including latency, body curvature, angular velocity, and duration (Burgess and Granato 2007b). We found that 2,4-DAB at 500μM, but not BMAA or AEG, increased the curvature of the C-bend response, as shown in three representative examples (**Figure 5A**) and quantified in **Figure 5C**. We also observed that the average latency of response in fish exposed to BMAA, and AEG was significantly reduced (**Figure 5B**). The BMAA/AEG binary mixture produced the same effect on latency as the individual toxins (**Supplemental Figure 2A**), indicating that at 250μM BMAA and AEG interact in an additive manner *in vivo* to alter startle performance (**Supplemental Figure 2A**). We used the same axial simplex design and statistical model to estimate the main effects and toxin interactions across the seven mixture ratios. The statistical significance and effect values were both confirmed for latency and curvature responses using the same linear model described above (**Figure 5D-E; Tables 4-5**), indicating that BMAA and AEG drive the decrease in latency and that 2,4-DAB alone drives the increase in curvature. Consistent with the increase in curvature we observed, the 2,4-DAB-exposed larvae also displayed higher angular velocity and longer duration of the SLC response (**Figure 5F-G**). Together, our behavioral analyses show that AEG affects larval zebrafish behavior similarly to BMAA, while 2,4-DAB alters distinct aspects of behavior from the other two toxins, playing a critical role in regulating coordinated movements in the zebrafish larvae model.

**Table 4.**
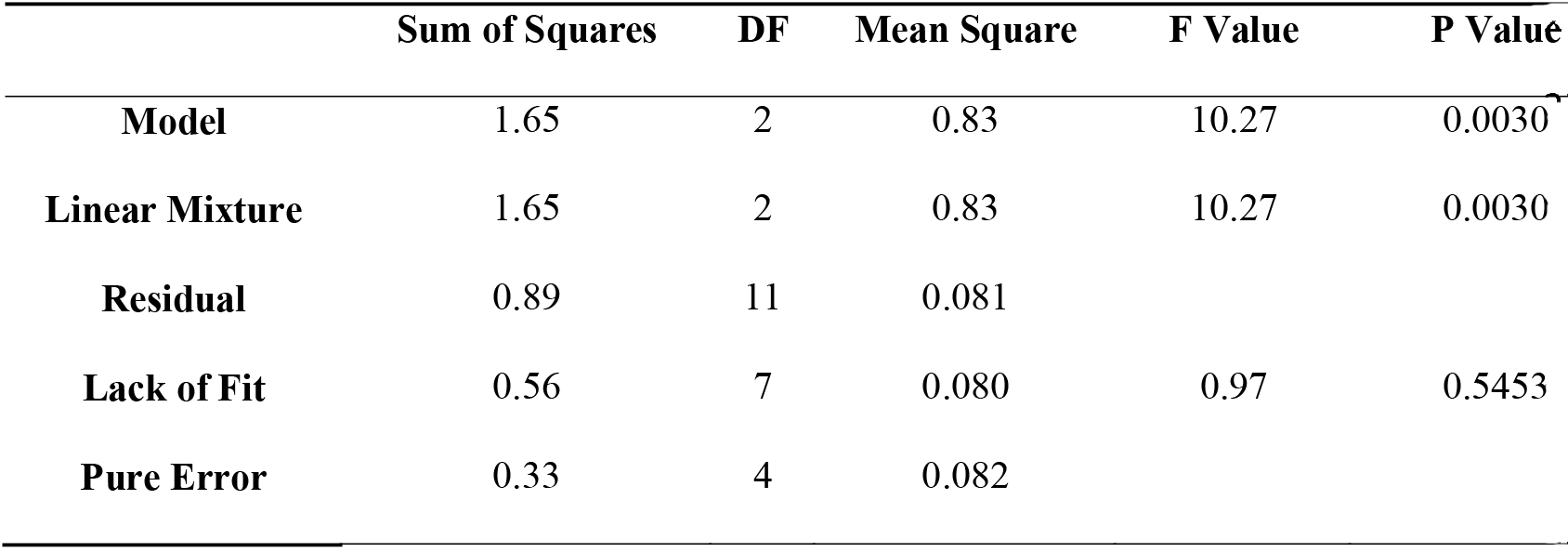
Analysis of Variance (ANOVA): Latency. ANOVA results, showing the significance a linear regression model (DF – degrees of freedom).

**Table 5.**
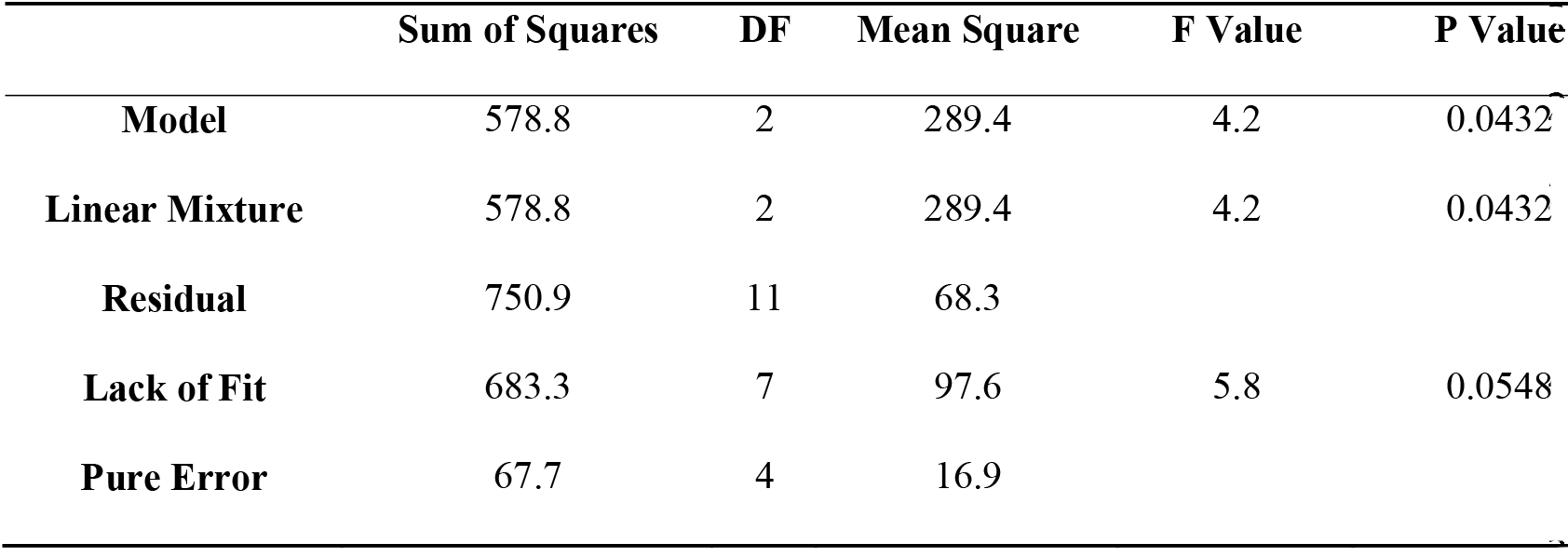
Analysis of Variance (ANOVA): Curvature. ANOVA results, showing the significance a linear regression model (DF – degrees of freedom).

**Figure 5.**
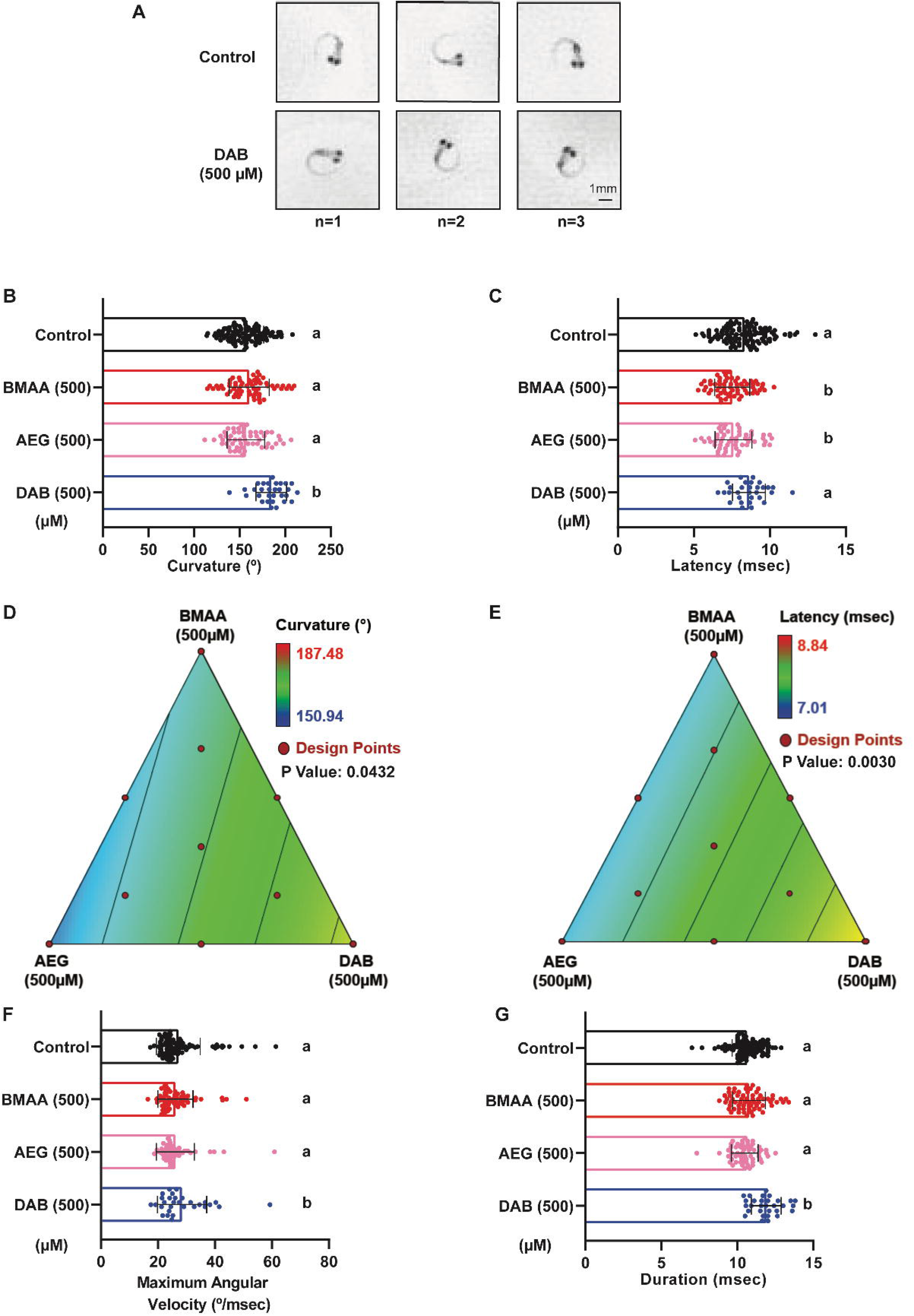
2,4-DAB modulates kinematics of the C startle response. (A) Three representative images of the peak SLC curvature in 6 days post fertilization (dpf) larvae exposed to vehicle (control, top) and to 2,4-DAB at 500μM (bottom). (B and C) Bar graphs quantifying the responses latency and curvature, respectively. (D and E) 2D contour plot for both the latency and curvature responses at different points in the design space, including each individual cyanotoxin (BMAA, AEG and 2,4-DAB) and their seven different mixture ratios (Linear regression model, P<0.05). (F and G) Bar graphs quantify responses maximum angular velocity and duration, respectively. Levels not connected by the same letter are significantly different (Tukey-Kramer HSD, Alpha 0.05).

### 3.5 Exposure to 2,4-DAB affects biological processes and protein homeostasis in vivo

Our data show that 2,4-DAB not only alters motor function during the startle response but more significantly reduces viability of larval zebrafish by approximately 50% by 4 days of development (**Figure 2A**). To understand the molecular processes by which 2,4-DAB may exert these effects we used a “bottom-up” shotgun proteomics approach. To identify both the mechanisms that may cause death as well as those that may induce behavioral neurotoxicity, we collected larval zebrafish exposed to 500μM 2,4-DAB at two time points: 2 dpf, prior to most of the toxin-induced mortality, and 4 dpf, after the wave of mortality had passed (**Figure 2A**). Control and 2,4-DAB-exposed larvae were each pooled in three biological replicates and flash frozen in liquid nitrogen, followed by proteomic analysis via LC-MS/MS. Approximately 2200 proteins were identified in each sample.

First, we analyzed the regulation of protein abundance by identifying all differentially expressed proteins (DEPs) in the 2,4-DAB samples by comparing the mean abundance at each time point (i.e., 2 dpf and 4 dpf) to their respective controls using a two-way Student’s t-test (P<0.05). All DEPs in both larval groups along with their significance values and fold change can be found in **Supplemental Table 1**, and volcano plots illustrating the direction of protein regulation at both time points are shown in **Supplemental Figure 3**. Overall, changes in protein levels at 2 dpf were less pronounced than those at 4 dpf. A total of 102 and 398 differentiated expressed proteins (DEPs) were identified for the 2 dpf and 4 dpf groups, respectively, representing a 4-fold increase in the number of dysregulated proteins in the 4 dpf samples (**Figure 6A**). There were no DEPs that were dysregulated at both time points, as illustrated by the absence of overlap in the Venn diagram in **Figure 6A**. These results suggests that 2,4-DAB affects an entirely different set of molecules during early embryonic development than at later larval stages.

**Figure 6.**
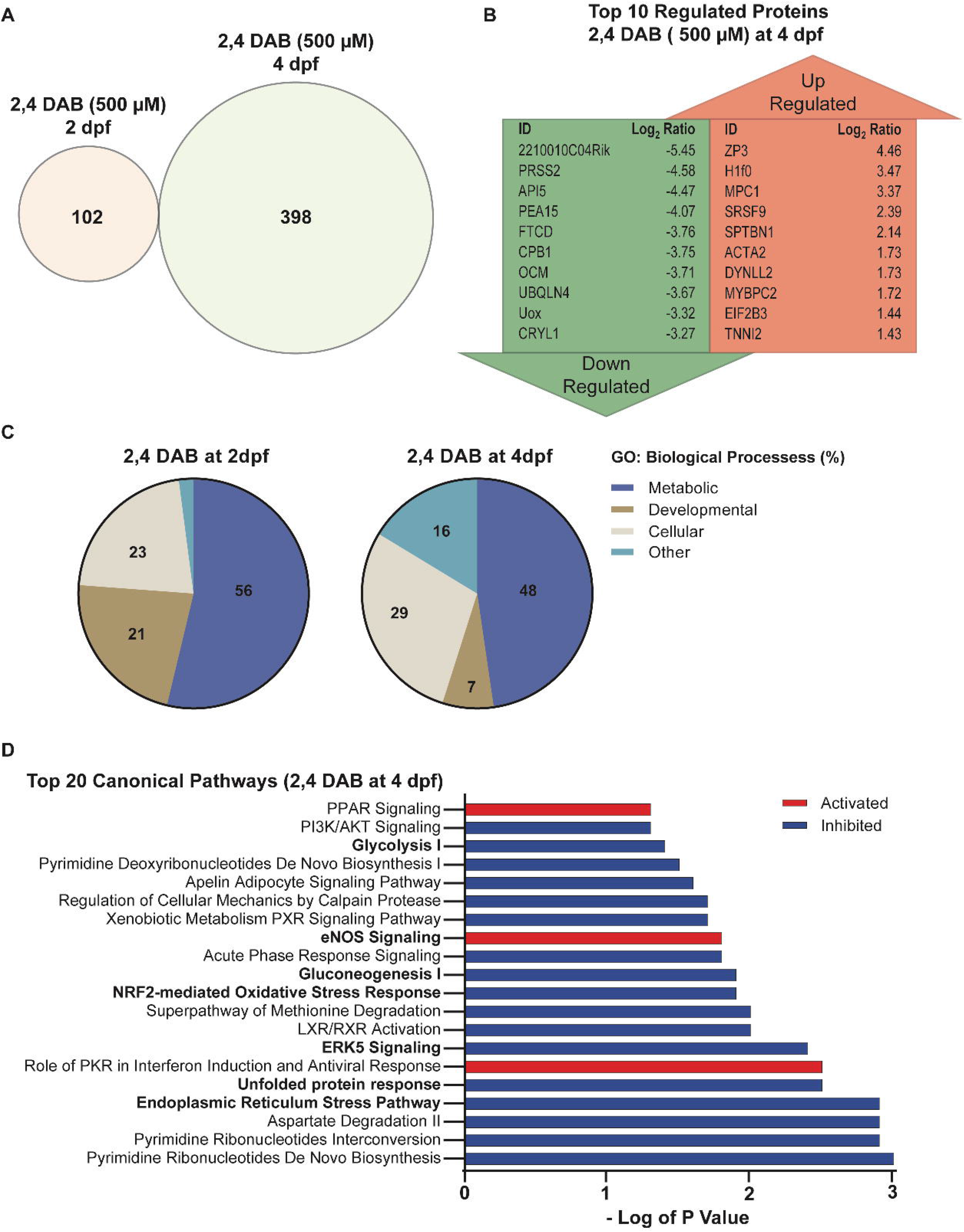
(A) Venn diagram shows no overlap in the differentially Expressed Proteins (DEPs) between 2,4-DAB (500μM) zebrafish larvae exposed groups at 2 dpf (left) and 4 dpf (right). (B) Top 10 DEPs found to be up or downregulated in the zebrafish larvae groups that were exposed to 2,4-DAB from 6 hpf until 4 days post fertilization (4 dpf). (C) Pie chart showing the percentage of enriched GO Biological Processes derived from each set DEPs time points: 2- and 4-day post fertilization. (D) Top 20 canonical pathways found to be activated or inhibited in the zebrafish larvae groups that were exposed to 2,4-DAB until 4 days post fertilization (4 dpf). Red shading indicates predicted activation (z score > 2), and blue shading indicates predicted inhibition (z score <-2). This bar graph width represents enrichment values in the form of minus log of p value that were greater than 1.3.

The top 10 up and down regulated proteins in the 4 dpf group (**Figure 6B**) included several associated with apoptosis (API5, PEA15) and metabolic function (MPC1, FTCD, UOX). To determine whether these functions were significantly affected by 2,4-DAB at both timepoints, we performed gene ontology (GO) enrichment analysis to reveal the classification of the biological processes represented by all DEPs. DEPs were classified into three major functional groups, including metabolism, development, and cellular functions. As shown in **Figure 6C**, both time points reveal the dysregulated proteins are largely involved in metabolic processes (2 dpf: 56%; 4 dpf: 48%), cellular processes (2 dpf: 23%; 4 dpf: 29%) and developmental processes (2 dpf: 21%; 4 dpf: 7%). A list of all subgroups of impacted biological processes can be found in **Supplemental Tables 3-4**. We next used Ingenuity Pathway Analysis (IPA) software to identify perturbed canonical pathways following exposure to 2,4-DAB. Canonical pathways were predicted using a Fisher’s exact t-test to determine the probability that DEPs from the dataset correspond with targets which are known to be activated/inhibited by those molecules based on knowledge in the Ingenuity database (Krämer et al. 2014). Only two canonical pathways were significantly enriched at 2 dpf: 1) neuronal nitrous oxide synthetase (nNOS) signaling and 2) protein ubiquitination pathway (**Supplemental Figure 4A**). Moreover, only eight proteins were significantly dysregulated within these canonical pathways in our samples (**Supplemental Figure 4B**), but both canonical pathways are associated with neuropathology (Atkin and Paulson 2014; Yi et al. 2009). We identified 56 canonical pathways that were significantly impacted in the 4 dpf samples (**Supplemental Table 2**). **Figure 6D** shows the top 20 canonical pathways that were predicted to be altered by 2,4-DAB (500μM) exposure, based on a p-value less than 0.05 and an activation z-score that was greater than an absolute value of 2. In agreement with our GO analysis, several of these canonical pathways are involved in cellular processes such as energy metabolism and protein homeostasis. For instance, both gluconeogenesis and glycolysis pathways were predicted to be inhibited (z-score<-2), leading to the predicted activation of metabolic diseases such as glucose metabolism disorders (z-score=2.227; overlap p-value=1.46E-03). Moreover, five key regulators in the data set (HSPA5, CAT, SOD2, HSP90, SOD1) were altered in the direction consistent with increased protein damage (z-score=2.219; overlap p-value=2.67E-04). These proteins are known to decrease damage of proteins in neurons (Mattson 2006; Murakami et al. 2011), and they were all found to be downregulated in our samples.

During normal development from 2 to 4 dpf, widespread changes in gene expression occur. To determine how 2,4-DAB exposure affects this developmental program, we identified DEPs between the 2 and 4 dpf control group and compared this set of proteins to the DEPs that were found between the 2 and 4 dpf 2,4-DAB exposed group. This systematic analysis revealed 868 DEPs for the control group and 606 DEPs for the 2,4-DAB-exposed group, with 349 DEPs common to both groups (**Figure 7A**). We found that DEPs in the control group were mostly upregulated while those in the 2,4-DAB-exposed group were mostly downregulated, perhaps reflecting a widespread impairment in the ability to activate gene expression (**Supplemental Figure 5**). Our enrichment analysis of control and 2,4-DAB DEPs revealed 7 highly enriched canonical pathways (overlap p-value<0.05 and z score>|2|) (**Figure 7B**). In agreement with our previous enrichment analysis showing downregulation at 4 dpf (**Figure 6C**), glycolysis and gluconeogenesis were both strongly activated in the control samples but only mildly activated in the 2,4-DAB samples. Protein kinase A and 14-3-3 protein signaling were similarly impaired in their activation by 2,4-DAB relative to control, while calcium signaling was more strongly activated by 2,4-DAB than in controls.

**Figure 7.**
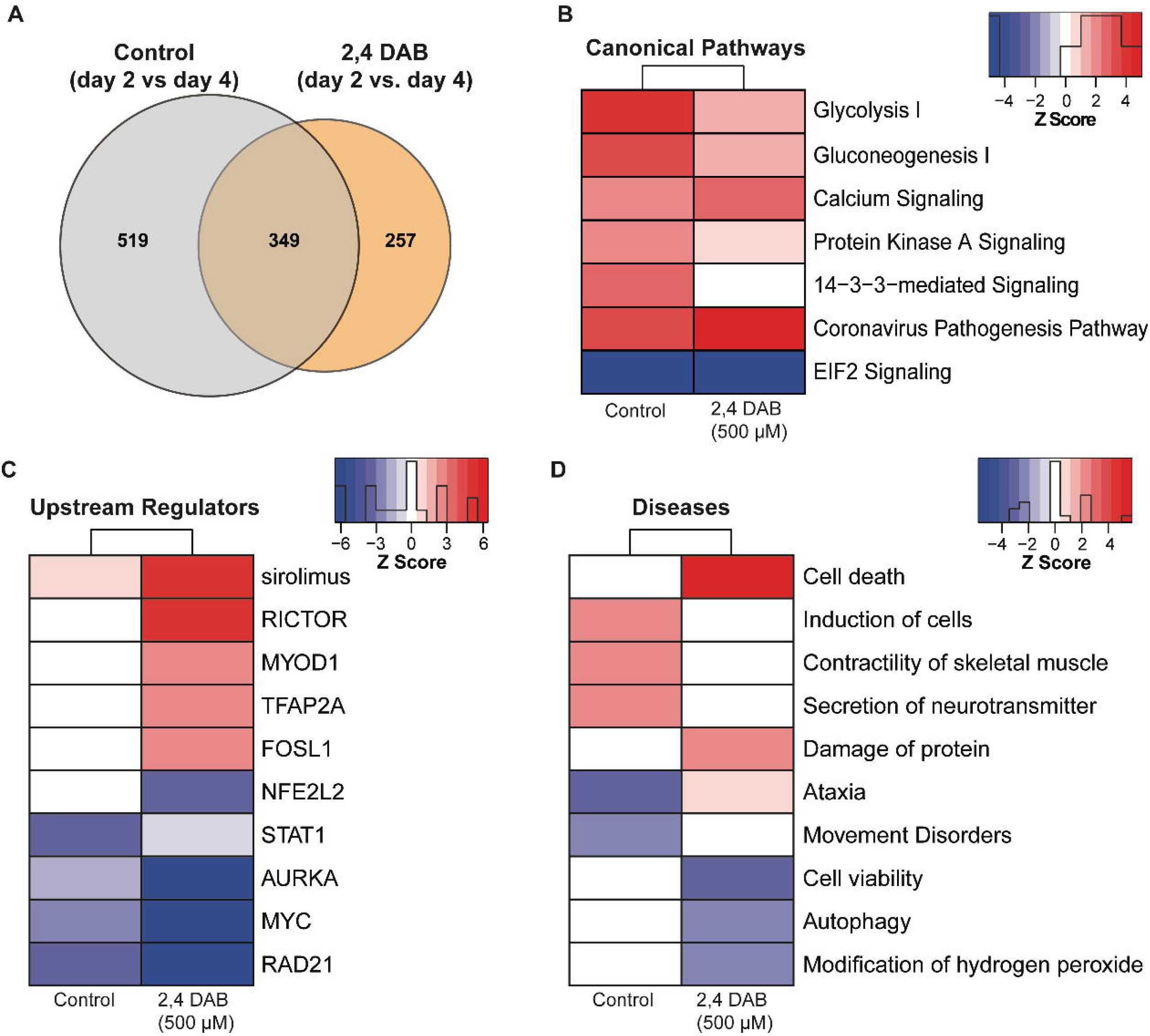
Systematic proteomics analysis of developmental changes in protein expression in control and 2,4-DAB (500μM) treated zebrafish larvae. (A) DEPs were identified in by comparing 4 dpf and 2 dpf samples in control and 2,4-DAB-treated larvae. Venn diagram showing DEP overlap between control and the 2,4-DAB treatment. (B) Canonical pathways impacted during development from 2 to 4 dpf in control and 2,4-DAB samples. (C) Predicted upstream regulators (D) Diseases and biological functions. The red or blue colored rectangles in each column indicates the z-score activities for each analysis. Red shading indicates predicted activation and blue shading indicates predicted inhibition. This heat map represents the activated of inhibited z-scores that were greater them the absolute value of 2.

To further investigate how 2,4-DAB impacts developmental protein regulation, we next performed upstream regulation and disease prediction analyses of the control and 2,4-DAB DEPs. The upstream regulator analysis evaluates linkage to DEPs through coordinated expression. Thus, it identifies potential upstream regulators (i.e., transcription factors or signaling proteins) that have been observed experimentally to affect protein expression (Krämer et al. 2014). Here, we have presented a subset of the top scored (overlap p-value<0.05 and z score>|2|) upstream regulators and diseases (**Figure 7C-D**). All activated and inhibited upstream regulators can be found in **Supplemental Table 7**. Among the top upstream regulators, we found that RICTOR was aberrantly strongly activated in the 2,4-DAB samples (z score= 5.18, **Figure 7C**) but not the control samples. RICTOR is a key component of mTOR signaling, which was highly enriched in our samples (p-value= 0.0007; **Supplemental Table 6**) and regulates cell proliferation and neurodevelopment (Karalis and Bateup 2021). Two additional upstream regulators and known transcription factors, NFE2L1 and 2, were predicted to be inhibited (overlap z score=−2.83) in the 2,4-DAB exposed samples (**Figure 7C**). NFE2L1 and NFE2L2 are involved in regulation of several downstream molecules involved in protective responses due to injury and inflammation (Saha et al. 2020). In addition to these regulators, multiple other neurodegeneration-related factors were identified (**Figure 7C; supplemental table 6**). Several diseases and biofunctions that implicate motor control were affected by the changes in protein expression from 2 to 4 dpf in control and 2,4-DAB-exposed fish (**Figure 7D**). Control samples showed an increase in muscle contractility and neurotransmitter secretion that was absent from the 2,4-DAB samples, and controls had decreased association with ataxia and movement disorders that was not seen in 2,4-DAB. These changes suggest reduced motor function in the 2,4-DAB-exposed larvae, which is consistent with the altered startle kinematics we observed in our behavioral analysis (**Figure 5**).

Finally, we explored the effects of 2,4-DAB on biological networks in zebrafish larvae. We found significant differential regulation of key proteins associated with cellular damage and inflammatory response (**Supplemental Table 5**). Within the highest scored network (score=68), we found that the ALS-associated superoxide dismutase 1 (SOD1; Enrichment p-value= 0.012; Log_2_ fold-change= −1.41) was significantly downregulated in our samples. We also found that another ALS-associated protein, ubiquilin 4 (Brettschneider et al. 2012; Edens et al. 2017), was one of the top 10 most downregulated proteins in the 4 dpf 2,4-DAB samples (UBQLN4; Enrichment p-value= 0.012; Log_2_ fold-change= −3.67; **Figure 6B**). These data demonstrate the power of proteomic screens to identify developmental effects of toxin exposures. Furthermore, this study indicates that 2,4-DAB impacts neurodegenerative pathways in larval zebrafish, highlighting the importance of further exploring the links between BMAA isomers and neural disease.

## 4. Discussion

Here, we provide a detailed report of the *in vivo* toxic effects of the beta-n-methylamino-l-alanine (BMAA) isomer 2,4-diaminobutyric acid (2,4-DAB) on zebrafish larvae. For many years, BMAA has been studied for its association with sporadic neurodegenerative disease, with several epidemiological studies revealing clusters of cases linked with environmental exposure to BMAA (Caller et al. 2009; Field et al. 2013; Masseret et al. 2013). However, BMAA is produced by cyanobacteria alongside its structural isomers 2,4-DAB and N-(2-aminoethyl) glycine (AEG) in natural algal blooms (Jungblut et al. 2018; Lance et al. 2018; Vo Duy et al. 2019). Although BMAA is a well□established neurotoxin, the toxicity of its isomers is less clear. Some recent *in vitro* studies have found that AEG and 2,4-DAB are toxic to neurons in culture (Main et al. 2018; Martin et al. 2019), but whether they exert neurotoxic effects *in vivo* has not been established. To the best of our knowledge, only one prior study has tested the toxic effects of 2,4-DAB in animals (Ferraiuolo et al. 2019), while none have evaluated the toxicity of AEG in an animal model or examined whether these cyanotoxins interact. Thus, we aimed to measure both the individual and the combined toxic effects of BMAA, AEG, and 2,4-DAB *in vivo*. Using the larval zebrafish model, we identified impacts on organismal viability, sensorimotor behavior, and global protein expression after developmental exposure to 2,4-DAB.

Our data show that 2,4-DAB is a more potent toxin than BMAA and AEG *in vivo*. The principle finding from our study is that 2,4-DAB caused 50% mortality at 500μM after just 4 days of exposure, while BMAA and AEG caused very little mortality under the same conditions (**Figure 2**). The effects of 2,4-DAB on viability are dose dependent, as the lowest observed effect level was 250μM (data not shown). These results are supported by a previous dose response study using zebrafish larvae in which 300μM was sufficient to produce signs of cardiotoxicity (Ferraiuolo et al. 2019). These results contrast with those from *in vitro* studies in which 2,4-DAB did not elicit an effect on viability at concentrations lower than 2000μM or 1000μM in human liver/ brain cancer cells or mouse motoneuron-like cells, respectively (Ferraiuolo et al. 2019; Martin et al. 2019). This distinction underscores the importance of studying toxin exposures in an *in vivo* context. Although exposure to BMAA or 2,4-DAB alone has previously been shown to cause morphological defects in larval zebrafish (Ferraiuolo et al. 2019; Purdie et al. 2009), we did not observe any significant morphological defects upon exposure to BMAA, AEG, or 2,4-DAB at 500μM. This could be due to differences in zebrafish strain, embryo medium, exposure route, or observer analysis. However, the lack of overt defects in the present study is consistent with our previous report on BMAA and Microcystin LR (MCLR) exposure (Martin et al. 2020).

Real-world cyanotoxin exposures are likely to involve more than one toxin, and our previous studies indicate that BMAA, AEG, and 2,4-DAB can interact *in vitro* (Martin et al. 2019), and that BMAA and MCLR can interact *in vivo* (Martin et al 2021). Building on these findings, here we employed a simplex axial design to investigate the combined effects of exposure to BMAA, AEG, and 2,4-DAB *in vivo*. Although Design of Experiments (DOE) approaches are typically used to determine the optimum combination of constituents that generate an outcome (e.g., for drug formulations) (N. Politis et al. 2017), several studies have shown that DOE is also an excellent tool for toxicological investigations of mixtures (Groten et al. 2001; Pomati et al. 2008). Based on our results using the simplex axial design, we suggest evaluating multiple endpoints (e.g., viability, spontaneous movement, startle kinematics, etc.) to further establish the significance of observed effects. Even though BMAA, AEG, and 2,4-DAB did not cause substantial effects on spontaneous locomotion on their own, exposure to the binary mixtures resulted in a significant modulation, decreasing both total distance travelled and average speed (**Figure 3**). Our spontaneous movement data also show that while AEG only exerts a minor effect on viability of larval zebrafish, it still can induce neurotoxicity at a behavioral level, and that that effect was enhanced when AEG was combined with 2,4-DAB (**Figure 3**).

Our analysis of sensorimotor function using an acoustic startle assay with precise kinematic tracking further revealed that BMAA, AEG, and 2,4-DAB all impair startle performance. We have previously shown that BMAA increases the frequency of Mauthner-cell dependent short-latency C-bends (SLCs) in larval zebrafish (Martin et al 2021), and our current data reinforce that finding (**Figure 4A**), indicating that the startle circuit is made hyperexcitable by BMAA. The isomer AEG also elicited the same hypersensitivity phenotype, and both BMAA and AEG decreased startle latency, suggesting they may act through similar mechanisms. Supporting this possibility, we also found that the binary mixture of BMAA and AEG produced the same hypersensitivity and latency phenotypes **(Figure 4B, Supplementary Figure 2A**), indicating that they likely interact in an additive manner. AEG has been shown to activate metabotropic glutamate receptor 5 (mGluR5) (Schneider et al. 2020), and BMAA can also act through mGluRs as well as NMDA receptors (Chiu et al. 2012; Pierozan and Karlsson 2019). Thus, there is strong evidence that both BMAA and AEG exert their neurotoxic effects by agonizing glutamate receptors.

Although 2,4-DAB can also agonize NMDA receptors and depolarize neurons (Spasic et al. 2018), we did not observe the same hypersensitive startle phenotype in 2,4-DAB-exposed zebrafish larvae. Rather, we observed that 2,4-DAB modulates startle kinematics, suggesting that 2,4-DAB acts through distinct mechanisms to cause neurotoxicity. The performance of SLC responses, defined through kinematic parameters such as curvature, angular velocity, and duration, depends on the proper function of the underlying motor circuits in the hindbrain and spinal cord, along with that of axial muscles. If startle kinematic defects are observed, this is an indication of motor dysfunction (Spasic et al. 2018). As 2,4-DAB did not alter the frequency or latency of SLCs, in contrast to BMAA and AEG, but instead impacted curvature, angular velocity, and duration (**Figure 5**), 2,4-DAB likely acts downstream of the command-like Mauthner neurons. In a recent forward genetic screen in larval zebrafish, several genes were identified that regulate the magnitude of the startle response, with mutations in *prdm12b*, *dolk*, and *kcna1a* causing the same increase in body curvature (Meserve et al. 2021), in which it was seen in our 2,4-DAB-exposed larvae. Dolichol kinase (dolk), a key enzyme in the protein glycosylation pathway, can regulate the hindbrain expression of the potassium Shaker-like channel encoded by *kcna1a* (Meserve et al. 2021), and mutations in *dolk* have been identified in patients with neuropathological conditions such as seizures (Helander et al. 2013) and autism spectrum disorder (Xiong et al. 2019). These studies support the idea that 2,4-DAB impacts similar molecular pathways to cause neurotoxicity.

Our unbiased proteomics analysis following *in vivo* 2,4-DAB exposure, to the best of our knowledge the first such dataset, further implicates 2,4-DAB in neurotoxic processes. We found that at both 2 dpf and 4 dpf 2,4-DAB strongly impacts metabolic function (Figure 6C), and consistent with the exaggerated startle phenotype caused by 2,4-DAB and shared with *dolk* mutants, 2,4-DAB inhibited glycosylation (**Figure 6D, 7B**). We also found that 2,4-DAB strongly disrupts several canonical pathways related to protein damage, including those associated with the NRF2-mediated oxidative stress response, the unfolded protein response, and endoplasmic reticulum stress (**Figure 6D**). These pathways have been implicated in neurodegeneration, which often features an increased incidence of misfolded proteins within neurons (Zhang et al. 2018). Interestingly, disruption of molecules involved in these pathways has also been observed in BMAA-exposed zebrafish (Frøyset et al. 2016). Moreover, the NRF2 oxidative stress response and the nitric oxide synthase (NOS) signaling pathways play important roles in maintaining cellular redox homeostasis and are implicated in ALS pathogenesis (Steinert et al. 2010; Tripathi et al. 2020). We found that 2,4-DAB-induced activation of the NOS signaling pathway was significant after just 2 days of exposure and was further enhanced with continued exposure at 4 days. Disruption of these pathways indicates the likelihood of protein damage due to 2,4-DAB exposure. For example, the nitric oxide radical is a known intracellular secondary messenger that can also react with superoxide anions to form peroxynitrite, which is a highly reactive molecule that can cause irreversible cell damage (Böhme et al. 1993). Consistent with our results, a previous study found that downregulation of NRF2 promotes increased levels of redox species and reduced expression of SODn1, a NRF2-dependent gene (Zou et al. 2016). We also observed significant downregulation of wild-type SOD1 and NFE2L2, and both SOD1 and NFE2L2 have been strongly associated with neurodegeneration (Baskoylu et al. 2018; von Otter et al. 2010). Further supporting a link between 2,4-DAB and motor neuron disease, the ALS-associated gene Ubiquilin 4 (UBQLN4) was among the top ten most strongly downregulated proteins in our 2,4-DAB-exposed samples. Ubiquilin 4 regulates autophagy and protein degradation, and expression of an ALS patient-derived gene variant in mouse motor neurons and zebrafish altered motor axon morphology (Edens et al. 2017). Thus, the molecular perturbations induced by 2,4-DAB exposure align with our behavior results and are consistent with neurodegenerative processes.

## 5. Conclusions

Altogether, our findings provide new evidence that 2,4-DAB drives toxicity in zebrafish larvae. Although all three cyanotoxins shows signs of both toxicity and neurotoxicity in larval zebrafish, 2,4-DAB causes both increased mortality and motor dysfunction. Finally, developmental exposure to 2,4-DAB affects zebrafish protein homeostasis and biological processes related to neurodegeneration. These results highlight the need for further investigation into 2,4-DAB’s role in disease and the specific mechanisms by which it exerts its effects.

## 6. Acknowledgement

We are thankful for startup funds provided by North Carolina State University (NCSU) and for pilot project support from the Center for Human Health and Environment (P30 ES025128). We are grateful to Derek Burton, MSc for zebrafish care and technical support and Dr. Jeffrey Enders, Ph.D. for feedback on the manuscript. The proteomics section of this work was performed using instrumentation from the Molecular Education, Technology and Research Innovation Center (METRIC) at NC State University.

## 7. Statements and Declarations

### Statement of Ethics

Animal experiments conform to internationally accepted standards and have been approved by the Institutional Animal Care and Use Committee (IACUC) at the University of North Carolina State University.

## 8. Abbreviations

SLC: short latency c-startle
LLC: long latency c-startle
HPLC: high-performance liquid chromatography
LC/MS: high-pressure liquid chromatography combined mass spectrometry
IPA: ingenuity pathway analysis
ANOVA: analysis of variance
GO: gene ontology
BMAA: β-methylamino-L-alanine
AEG: N-(2-aminoethyl)glycine
2,4-DAB: 2,4-diaminobutyric acid
DEPs: differentially expressed proteins

