## Supplemental Material for "The cyanotoxin 2,4-DAB enhances mortality and causes behavioral and molecular dysfunctions associated with neurodegeneration in larval zebrafish"

**^1^ Department of Biological Sciences, North Carolina State University, Raleigh, NC**

**Submitted to: Neurotoxicity Research**

**Submitted: 10/13/21**

**Pages: 24; Figures: 5; Tables: 7**

**Running Title: Investigation of the neurotoxicity of BMAA, AEG and 2,4 DAB in vivo.**

**Keywords: Cyanotoxins; Mixtures; 2,4 DAB; Zebrafish; Behavior; Proteomics**

***Author for Correspondence**

**Kurt C. Marsden**

**Department of Biological Sciences**

**North Carolina State University**

**Raleigh, NC**

****

**Supplemental Methods**

*Protein Extraction*

A pool of approximately 20 zebrafish larvae whole body for each condition were suspended in a 100 µL solution of 50 mM ammonium bicarbonate (pH 8.0) containing 1% SDC. Zebrafish bodies were lysed by probe sonication via 2 pulses at 20 seconds per pulse at a 20% amplitude setting. Cellular debris were removed via centrifugation at 10,000 RPM for 5 minutes. The supernatant was retained and assessed for protein quantification by bicinchoninic acid (BCA) assay.

*Protein digestion*

One hundred micrograms of protein were diluted to a final volume of 100 µL with the 50 mM ammonium bicarbonate and 1% SDC solution. DTT was added to a final concentration of 5 mM and incubated at 60ºC for 30 minutes to reduce disulfide bonds. Samples were cooled to room temperature, followed by alkylation of cysteine residues with iodoacetamide (15 mM) at room temperature and in the dark for 20 minutes. Samples were then subjected to filter-aided sample preparation (FASP) (Wisniewski et al. 2009) for protein cleanup using Vivacon 30,000 kDa molecular weight cutoff filters and reconstituted with 100 µL of 50 mM ammonium bicarbonate solution. Tryptic digestion was carried out by hydrating lyophilized trypsin to a stock solution of 1 µg/µL with 0.01% acetic acid in water followed by addition of trypsin to the protein mixture at a 1:50 ratio and then incubated at 37ºC for 4 hours. After digestion, peptides were acidified with HCl at a final concentration of 250 mM (pH ≤ 3).

*LC-MS/MS Analysis*

Three microliters of each sample were injected and analyzed using an Easy nanoLC 1200 coupled to an Orbitrap Exploris 480 Mass Spectrometer (Thermo Scientific, Bremen, Germany). LC separation was performed with an EASY-spray system, consisting of a 50 cm, 75 μm ID PepMap RSLC, C18, 100 Å, 2μm particles, in which was connected to an Easy-nLC Ultra UHPLC system (Thermo Scientific, San Jose, CA). Separation of peptides was achieved through a gradient of mobile phase A (98% water, 2% acetonitrile, and 0.1% formic acid) and mobile phase B (79.9% acetonitrile, 20% water and 0.1% formic acid). The LC method applied consisted of a gradient of 5%-50% B over 120 minutes, followed by a ramp to 95% B in two minute. The column was washed at 95% B for 16 minutes. Tandem mass spectrometry was carried out using positive ion mode and data dependent mode with a three second cycle time. MS1 scans were performed at a resolving power of 60,000 from m/z 375 to 1600 and an automatic gain control (AGC) target of 1 x 10^6. MS2 scans were performed at a resolving power of 15,000 and an AGC target of 5 x 10^4. A 30 second dynamic exclusion window was applied during sampling to avoid repeated MS2 interrogation of high-abundant species.

**Supplemental Figure Legends**

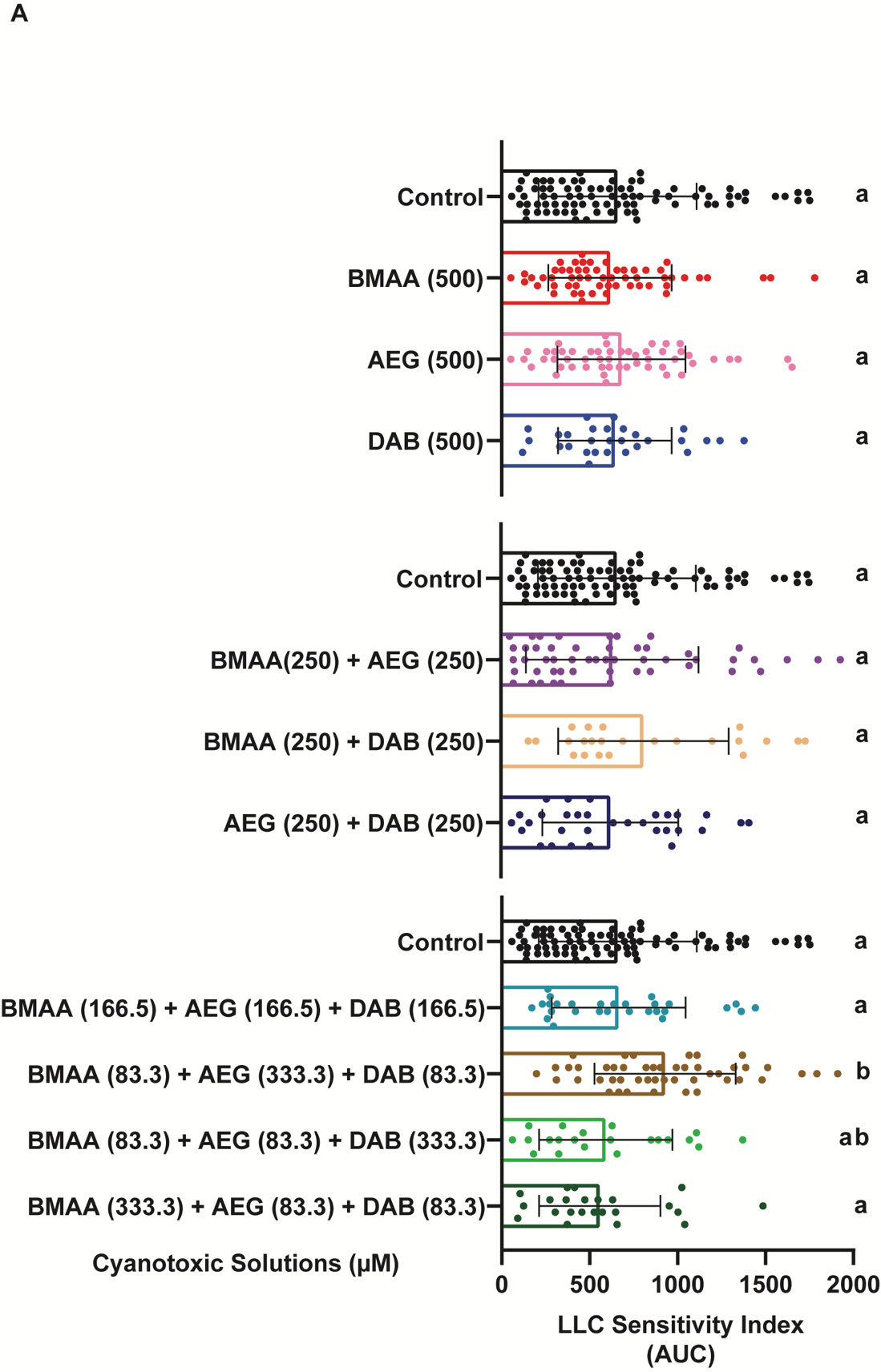

**Supplement Figure 1:** Long latency c startle (LLC) behavior is not affected by cyanotoxins. (A) Bar graphs display the distribution of the LLC sensitivity indices for each tested larva after exposure to individual cyanotoxins, binary mixtures and three component mixtures, respectively. n = 108 siblings; mean ± SEM. Levels not connected by the same letter are significantly different–Tukey-Kramer HSD, Alpha 0.05.

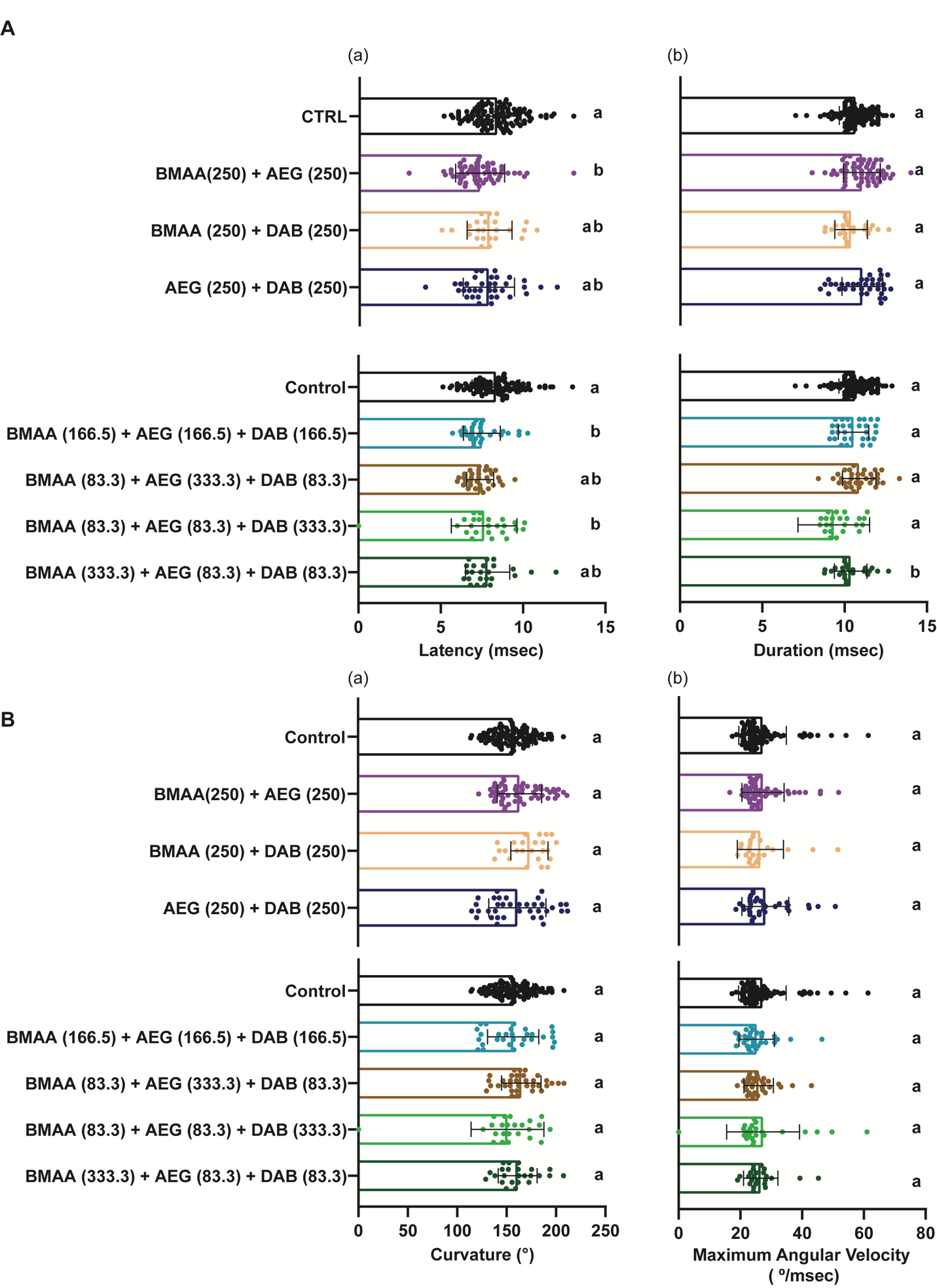

**Supplement Figure 2: Mixture of cyanotoxins does not affect larval zebrafish behavior.** (A) Bar graphs quantify responses latency and duration, respectively. (B) Bar graphs quantify responses curvature and maximum angular velocity, respectively. Analysis of head orientation allows for automated identification and measurements of kinematic responses of the c startle behavior. Levels not connected by the same letter are significantly different – Tukey-Kramer HSD, Alpha 0.05.

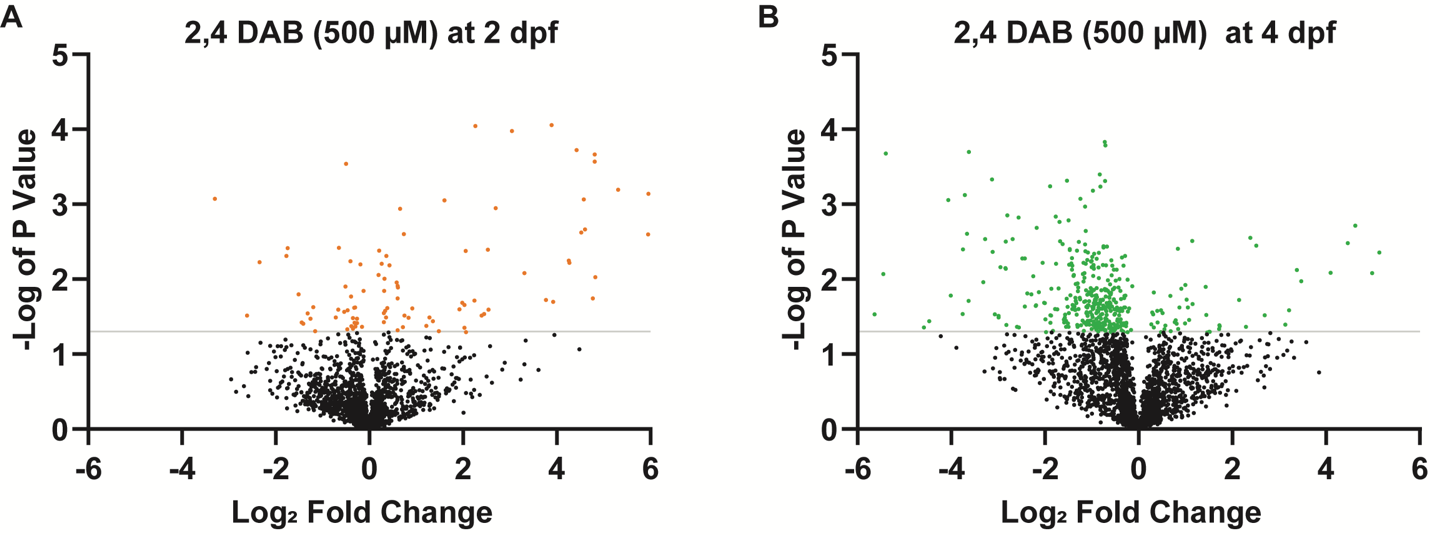

**Supplement Figure 3**. **Volcano plots**. (A-B) Volcano plots representation of regulated proteins. Orange and green dots represent significantly (p ≤ 0.05) regulated proteins for the 2 days post fertilization group and 4 days prost fertilization group, respectively. Black dot represents non-significantly (p ≥ 0.05) proteins compared to their respective controls.

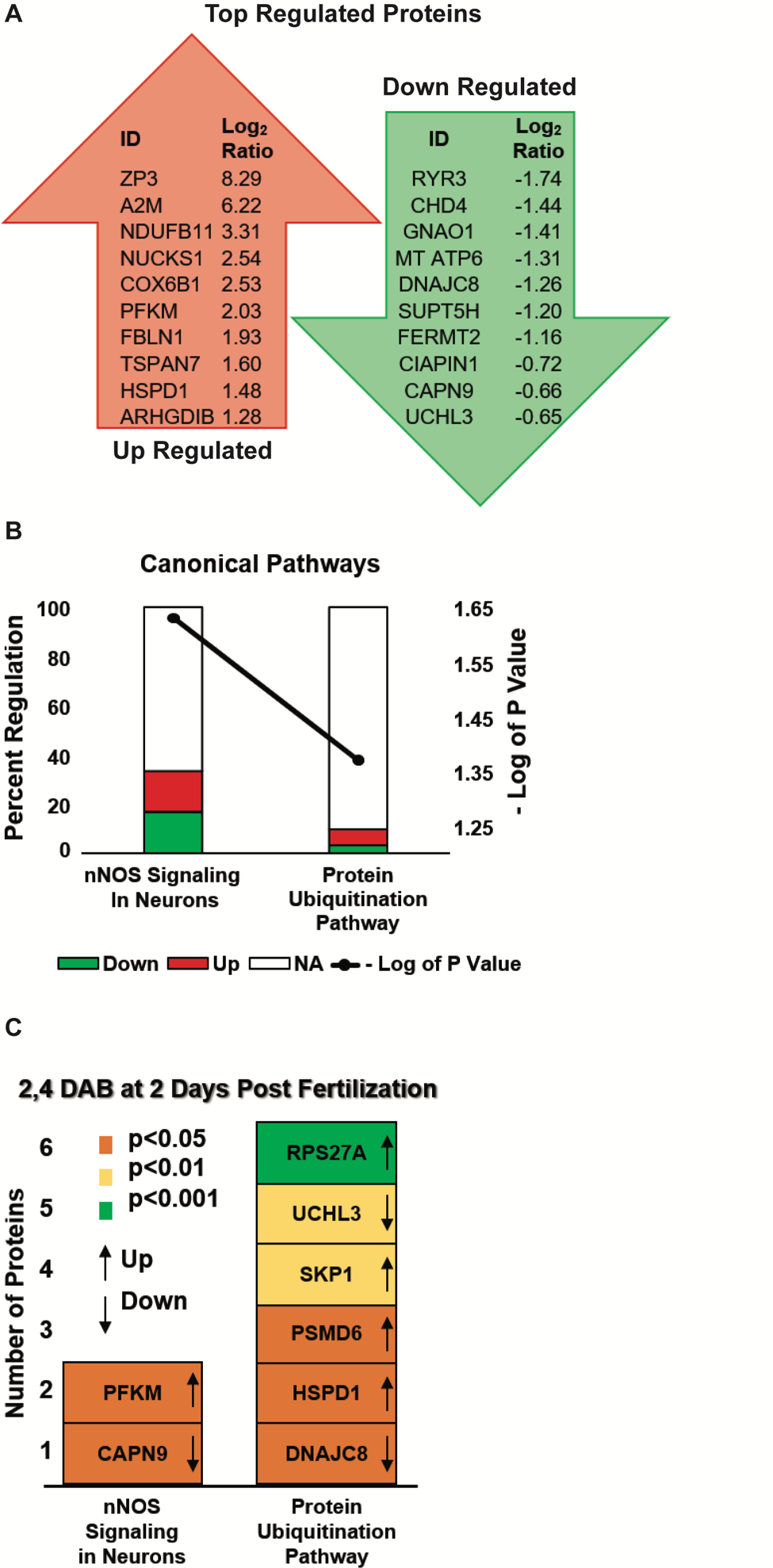

**Supplement Figure 4: 2,4 DAB at 500 µM causes minimal molecular changes at 2 days post fertilization.** (A) Top 10 differentially expressed proteins found to be up or downregulated in the zebrafish larvae groups that were exposed to 2,4 DAB until 2 days post fertilization (2 dpf). (B) Bar chart of the only two significantly enriched canonical pathways derived from the differentiated expressed proteins from the zebrafish larvae 2 dpf group. Left axis represents the percentage of DEPs found in pathway and right axis displays their significance in minus log of p value. (C) Bar graph showing regulation of enriched protein biomarkers followed by 2 days of 2,4 DAB 500 µM exposure.

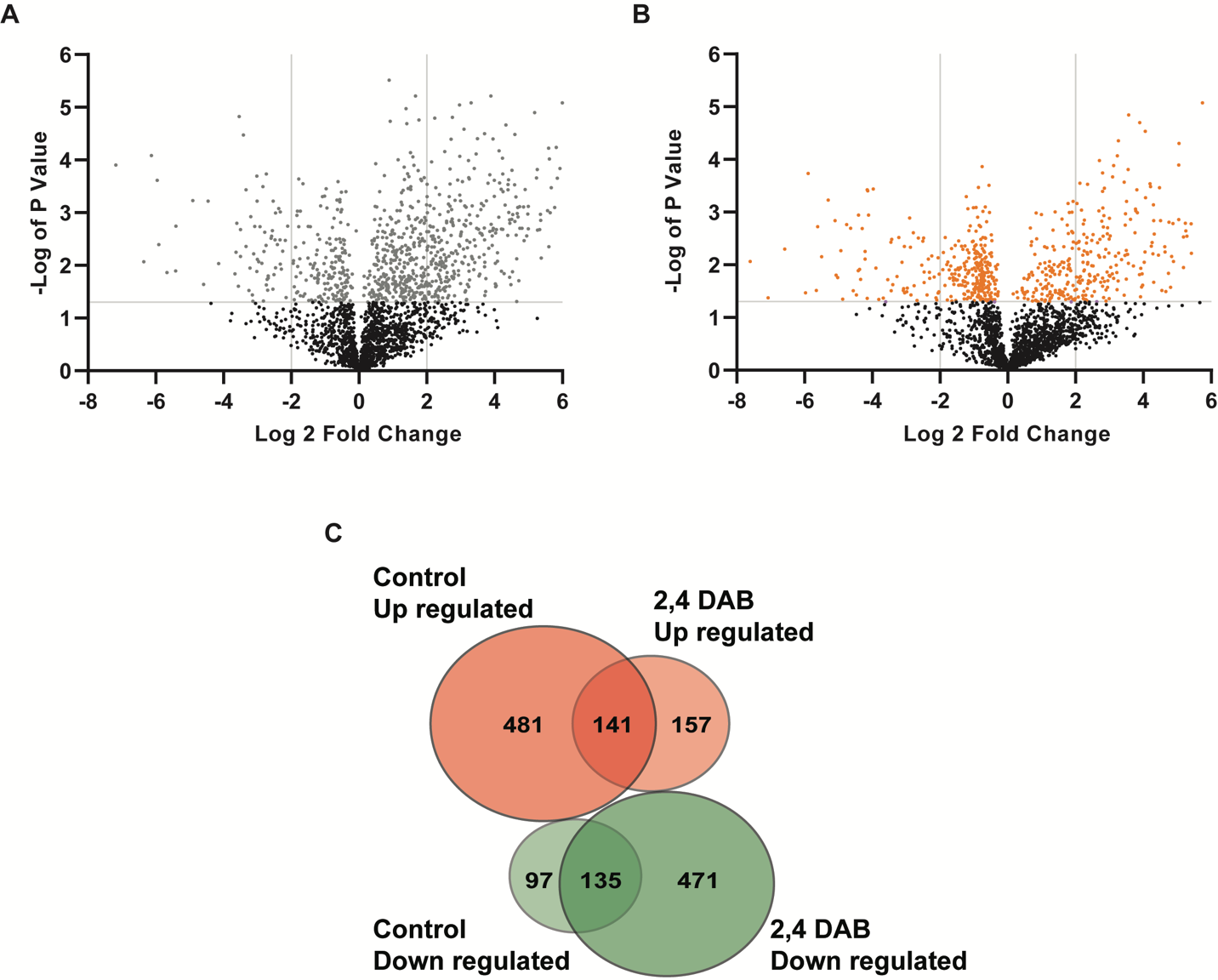

**Supplement Figure 5: Systematic analysis volcano plots**. (A-B) Volcano plots representation of regulated proteins. Gray and orange dots represent significantly (p value ≤ 0.05) regulated proteins for the control and 2,4 DAB exposed zebrafish larvae groups, respectively. Black dot represents non-significantly (p ≥ 0.05) proteins compared to their 2 days post fertilization time points. (C) Venn diagram showing protein overlap between control and the 2,4 DAB treatment. From top to bottom: Control and 2,4 DAB (500 µM) upregulated proteins (Red) and down regulated proteins (Green), respectively.

**Supplemental Tables and Legends**

| **2,4 DAB (500 µM) at 2 dpf** | | | **2,4 DAB (500 µM) at 4 dpf** | | |
| --- | --- | --- | --- | --- | --- |
| **T: Protein IDs** | **N: -Log Student's T-test p-value** | **N: Student's T-test Difference** | **T: Protein IDs** | **N: -Log Student's T-test p-value** | **N: Student's T-test Difference** |
| E9QHY5 | 4.0582 | 3.89242 | Q5RKP8 | 3.82992 | -0.723495 |
| A7MBZ3 | 4.04427 | 2.26146 | Q803I7 | 3.78745 | -0.712429 |
| A7MBZ5 | 3.98208 | 6.54447 | A2RV44 | 3.69622 | -3.62119 |
| A0A2R8RUH9 | 3.97736 | 3.04445 | A0A0R4IU03 | 3.67685 | -5.39335 |
| A0A286YAS1 | 3.93549 | 7.31167 | Q29RA4 | 3.39752 | -0.827574 |
| A5JET5 | 3.72475 | 4.4238 | Q7SY34 | 3.3326 | -3.13203 |
| A9JRE5 | 3.67158 | 7.43674 | F1QGK0 | 3.31447 | -1.52686 |
| F1QC84 | 3.66437 | 4.81155 | A0A2R8Q3A3 | 3.31055 | -0.716787 |
| E7F6Q4 | 3.56858 | 4.80839 | A9JRT0 | 3.23777 | -1.89038 |
| Q6PHF1 | 3.5393 | -0.499031 | B8A516 | 3.23665 | -0.814496 |
| Q1RM40 | 3.52488 | 6.80842 | Q6NZU0 | 3.18117 | -0.976518 |
| A8WFS6 | 3.36947 | 6.21725 | Q800A2 | 3.12301 | -3.70929 |
| Q6NW70 | 3.28577 | 8.2868 | Q66I80 | 3.07461 | -1.24447 |
| E7FCS3 | 3.19578 | 5.31067 | E7FEH9 | 3.05449 | -4.06614 |
| F1QY35 | 3.14101 | 5.95342 | Q1L8P1 | 2.97058 | -1.14025 |
| F1QVW8 | 3.07498 | 6.28033 | E9QHX7 | 2.85213 | -2.80204 |
| B3DFK4 | 3.07322 | -3.29978 | A0A0R4IKD6 | 2.83627 | -1.77165 |
| A0A2R8QN17 | 3.06482 | 4.57999 | A8KBV7 | 2.82468 | -2.56536 |
| Q642H0 | 3.05261 | 1.60278 | Q5SPD5 | 2.78379 | -1.49854 |
| H0WED7 | 2.94921 | 2.69402 | Q1JQ08 | 2.7633 | -1.69081 |
| Q7SXA3 | 2.94109 | 0.659412 | F1QUS3 | 2.71465 | 4.62 |
| A0A2R8RTZ6 | 2.73697 | 6.80017 | B2GS10 | 2.68689 | -2.17426 |
| E7FFZ3 | 2.66332 | 4.60283 | Q6P3J2 | 2.6447 | -1.13167 |
| F1RED5 | 2.6237 | 4.52534 | E9QFX2 | 2.6071 | -3.66567 |
| Q7T358 | 2.602 | 0.737941 | F1QVC6 | 2.55286 | 2.38661 |
| Q3B744 | 2.59649 | 5.94992 | Q0V973 | 2.53625 | -3.27424 |
| Q504C0 | 2.41874 | -0.648837 | Q5BLA2 | 2.53594 | -2.69125 |
| Q5RGQ4 | 2.41615 | -1.74401 | F1QNA7 | 2.5091 | 1.14347 |
| F1QIP9 | 2.39872 | 6.11185 | F8W5B2 | 2.50807 | -1.67826 |
| Q6DH63 | 2.39408 | 2.52792 | A0A0R4IAD4 | 2.50061 | -2.83958 |
| A0A0R4IKW3 | 2.38188 | 0.212849 | E9QDF8 | 2.49651 | -1.27598 |
| B2CQD3 | 2.37682 | 2.05888 | Q9PWC7 | 2.48179 | 4.46039 |
| Q4QRC7 | 2.30917 | -1.7705 | Q503E2 | 2.47047 | -1.62604 |
| Q6P959 | 2.30843 | 0.362307 | Q1LYC9 | 2.46271 | -1.14488 |
| F1REP2 | 2.24873 | 4.25418 | A7MBZ6 | 2.44695 | 2.50871 |
| Q6IQV5 | 2.23941 | -0.404558 | R4GEX2 | 2.43945 | -0.746715 |
| Q8QGV5 | 2.22758 | -2.34149 | F1R3J9 | 2.43669 | -0.675093 |
| B0V0J5 | 2.21699 | 4.27192 | Q6DI36 | 2.42372 | -0.751933 |
| Q6PC90 | 2.20791 | 0.261508 | A8KBR9 | 2.40667 | 0.838566 |
| Q6IQT1 | 2.1973 | -0.186917 | F1R5A5 | 2.40227 | -1.46816 |
| Q1ED17 | 2.18349 | 0.430036 | F1QJN9 | 2.39603 | -3.75071 |
| Q1LYN0 | 2.08027 | 3.31138 | A0A0R4IS05 | 2.38958 | -1.47175 |
| Q0D294 | 2.05553 | 0.197168 | F1QMC6 | 2.38272 | -1.31358 |
| A7E2J3 | 2.02884 | 4.82254 | A0A2R8RPQ9 | 2.37142 | -1.1395 |
| A0A0R4ID68 | 2.00771 | 0.321396 | Q3B7E4 | 2.36254 | -3.11588 |
| Q6PBY8 | 1.95546 | 0.580386 | Q9PW37 | 2.35571 | -1.08107 |
| Q6IQ64 | 1.91112 | 0.60131 | Q8JH37 | 2.35567 | 5.13687 |
| B3DFW4 | 1.90127 | -0.509134 | Q6NYA1 | 2.35368 | -0.849555 |
| Q5BJB9 | 1.88925 | 0.607092 | Q7SY03 | 2.33801 | -0.952874 |
| A0A2R8RHQ3 | 1.84465 | 0.31934 | A0A0R4IA10 | 2.31078 | -0.286714 |
| Q5TZ29 | 1.8422 | -0.126497 | E7FE90 | 2.29747 | -0.988392 |
| Q5XJ76 | 1.79724 | -1.5074 | Q7ZUJ8 | 2.28822 | -0.358571 |
| A0A0R4I9F5 | 1.76835 | -0.390299 | B8JL43 | 2.27947 | -1.42003 |
| F1RBW3 | 1.74323 | 0.603799 | Q7ZTW2 | 2.2771 | -2.48521 |
| A8BAS1 | 1.74273 | 4.76811 | Q08BF9 | 2.27699 | -2.43032 |
| A3KQR6 | 1.72362 | 3.76881 | Q568N9 | 2.27219 | -0.893655 |
| A9JRE3 | 1.71276 | 2.2415 | A2AR68 | 2.25614 | -1.09885 |
| F1QYE3 | 1.69603 | 3.92225 | A0A2R8QFM8 | 2.25255 | -0.586418 |
| B8A565 | 1.68571 | 1.98561 | F1QZN3 | 2.24268 | -1.00358 |
| X4YKC7 | 1.65454 | 2.03211 | Q8AYE3 | 2.23438 | -1.01782 |
| A0A0R4IBG8 | 1.62512 | -1.19725 | A0A0F6PK88 | 2.21883 | -1.17432 |
| F1R2K9 | 1.62391 | -0.303851 | Q6PC74 | 2.21841 | -2.05242 |
| Q8JH72 | 1.61695 | -0.324204 | F1QBW9 | 2.21034 | -1.43677 |
| B2GSF5 | 1.61434 | 0.381922 | Q1LYL6 | 2.20707 | -1.76517 |
| A0A0R4IDI3 | 1.61042 | 0.91652 | A9JRT8 | 2.20637 | -1.13116 |
| A0A0R4ILN6 | 1.59811 | 1.92649 | I3ITC7 | 2.19511 | -1.48446 |
| Q6DG44 | 1.59521 | -0.664821 | Q6PC65 | 2.1902 | -0.763195 |
| Q66I37 | 1.59349 | 2.54263 | Q6P5M5 | 2.18769 | -0.960421 |
| F1QET7 | 1.58419 | -0.463005 | E7FBD3 | 2.18486 | -0.696739 |
| Q1ED16 | 1.57734 | 0.336009 | Q6NYU8 | 2.18269 | -0.566284 |
| A0A0R4IY17 | 1.56562 | -0.534305 | A2BIM7 | 2.18071 | -1.41621 |
| Q8JHH3 | 1.54855 | 0.311961 | B3DKC0 | 2.18035 | -0.326256 |
| A0A0U2NKZ2 | 1.54182 | -1.31436 | A0A0J9YJC9 | 2.16095 | -1.00286 |
| Q1RMB1 | 1.53705 | 2.44674 | A0A2R8RJC4 | 2.16058 | -2.94922 |
| A8KB68 | 1.51572 | 2.39521 | Q6DRN5 | 2.15745 | -0.857063 |
| Q5BLE6 | 1.51519 | -2.60841 | F1Q7P7 | 2.15488 | -0.963346 |
| Q7ZWE5 | 1.51255 | 0.748178 | A7YT77 | 2.14848 | -2.84563 |
| B2GNT9 | 1.491 | -0.718457 | Q6PFT7 | 2.13998 | -0.95356 |
| Q6PC22 | 1.48838 | 0.359627 | F1R5W4 | 2.13914 | -2.83986 |
| Q7SZQ6 | 1.48772 | 1.28309 | A7YYE9 | 2.12636 | -0.528002 |
| Q5RH28 | 1.48695 | 0.834299 | Q803V8 | 2.1256 | -1.4253 |
| Q6IQI3 | 1.48129 | -0.351069 | Q6NYV1 | 2.12512 | -0.466992 |
| Q1ED27 | 1.4747 | -1.25903 | F1Q6Z3 | 2.12106 | 3.37445 |
| A7E2K4 | 1.47387 | -0.260997 | F1R4N4 | 2.08676 | 4.09859 |
| E7FDL8 | 1.43827 | 1.35618 | A9JSV9 | 2.08269 | 4.98599 |
| Q9I8U8 | 1.42887 | 0.307999 | Q6DHT4 | 2.07973 | -1.15372 |
| F1QWV5 | 1.42171 | -1.44157 | Q7SX88 | 2.06854 | -1.18948 |
| E7F0A1 | 1.41702 | -0.318904 | Q561Z7 | 2.06746 | -5.44761 |
| A0A0R4ICN9 | 1.41383 | -0.260391 | Q7ZW04 | 2.05776 | -1.38192 |
| F8W442 | 1.40752 | -1.41154 | Q503D5 | 2.05303 | -0.575359 |
| A0A0G2L5F9 | 1.37732 | 1.21986 | B3DFP9 | 2.04345 | -1.39274 |
| Q66HY3 | 1.37151 | -0.382655 | A0A0R4IM65 | 2.04278 | -0.967028 |
| A0A0R4IQN8 | 1.36678 | -0.298943 | B2BWH5 | 2.0402 | -2.24681 |
| A0A0R4IMS3 | 1.36408 | -0.155223 | Q1LXJ7 | 2.01816 | -1.15755 |
| Q7ZYX0 | 1.36046 | 0.718764 | A0A286YBA4 | 2.0164 | -0.850412 |
| Q66HV8 | 1.35004 | 2.0324 | F1QDQ1 | 2.00087 | -0.40832 |
| A0A0B5JWA3 | 1.34076 | -0.351802 | Q6P2M0 | 1.99565 | -0.23717 |
| Q7ZSZ0 | 1.33 | -0.471355 | H9GX78 | 1.98347 | -0.65601 |
| A4IGC2 | 1.31907 | 0.605777 | Q802C9 | 1.9825 | -0.493238 |
| Q803B0 | 1.30811 | 1.48428 | Q6NYV3 | 1.97237 | 3.47054 |
| A0A2R8RR15 | 1.30635 | -1.15719 | Q90Z38 | 1.9681 | -1.1735 |
|  |  |  | E9QFD2 | 1.96218 | -3.31978 |
|  |  |  | E7EZ90 | 1.94483 | -1.30019 |
|  |  |  | Q90Z37 | 1.93573 | -1.34261 |
|  |  |  | Q6NY25 | 1.92978 | -0.607272 |
|  |  |  | Q1L8Z0 | 1.92398 | 0.99628 |
|  |  |  | A2BGU3 | 1.90792 | -0.136616 |
|  |  |  | Q0D2W2 | 1.89896 | 1.4302 |
|  |  |  | Q7ZVJ4 | 1.89137 | -1.28913 |
|  |  |  | Q7SXK0 | 1.88891 | -0.431388 |
|  |  |  | Q803R9 | 1.8843 | -0.514395 |
|  |  |  | E7F4R9 | 1.88321 | -1.02264 |
|  |  |  | Q8AWD9 | 1.8822 | -0.895341 |
|  |  |  | Q6DH07 | 1.87788 | -1.7126 |
|  |  |  | F1QTE2 | 1.87475 | -1.86646 |
|  |  |  | Q6DRD6 | 1.87339 | 0.912005 |
|  |  |  | A3KNY9 | 1.86872 | -0.723478 |
|  |  |  | Q6NYJ5 | 1.86764 | -0.537466 |
|  |  |  | Q6TNV4 | 1.86294 | -0.55084 |
|  |  |  | Q24JW2 | 1.86101 | -0.687023 |
|  |  |  | Q7SXW9 | 1.86054 | -0.995616 |
|  |  |  | A0A0R4IHU9 | 1.85829 | -0.957825 |
|  |  |  | A0A2R8QTH3 | 1.85317 | -0.794828 |
|  |  |  | Q32PS5 | 1.85294 | -1.12349 |
|  |  |  | Q7ZVP2 | 1.85042 | -0.711229 |
|  |  |  | A9JT16 | 1.84529 | -0.657157 |
|  |  |  | E9QIF2 | 1.84323 | -0.392258 |
|  |  |  | Q6PC44 | 1.83764 | -1.29811 |
|  |  |  | A0A0B5JK63 | 1.82978 | -1.00081 |
|  |  |  | Q6DI22 | 1.82761 | -1.05468 |
|  |  |  | Q90Y38 | 1.82606 | -2.12686 |
|  |  |  | F6P9M5 | 1.82474 | 0.32185 |
|  |  |  | A0A2R8RKG2 | 1.82015 | -0.588513 |
|  |  |  | Q1LXD2 | 1.81972 | -0.621051 |
|  |  |  | B0V3S2 | 1.80926 | -2.37699 |
|  |  |  | A0A286YAP4 | 1.79989 | -0.593744 |
|  |  |  | A5PLF4 | 1.79896 | -0.606078 |
|  |  |  | Q6P124 | 1.79882 | -1.02623 |
|  |  |  | B0VEQ2 | 1.79713 | -2.29307 |
|  |  |  | Q1LXV5 | 1.79519 | -1.0979 |
|  |  |  | F8W3N3 | 1.78747 | -1.45636 |
|  |  |  | Q6DRE8 | 1.7837 | -0.906656 |
|  |  |  | A0A0G2KEX1 | 1.7824 | -0.994219 |
|  |  |  | F1QFK2 | 1.78014 | -0.92639 |
|  |  |  | F1QVD2 | 1.7798 | -0.285099 |
|  |  |  | A7MCJ2 | 1.77949 | -4.0131 |
|  |  |  | A0A2R8RPC7 | 1.7787 | 0.678196 |
|  |  |  | Q7T3B1 | 1.77513 | -0.300215 |
|  |  |  | Q568Q7 | 1.77044 | -0.702129 |
|  |  |  | F1Q9U6 | 1.76177 | -0.887945 |
|  |  |  | F1R1P6 | 1.76015 | -0.626989 |
|  |  |  | Q6NY60 | 1.75356 | -1.34856 |
|  |  |  | Q568H8 | 1.75303 | -0.622067 |
|  |  |  | I3ISH4 | 1.75057 | -1.2586 |
|  |  |  | Q29RA2 | 1.74697 | -0.309451 |
|  |  |  | A0A0N4SU18 | 1.7398 | -1.53683 |
|  |  |  | A0A2R8Q5K7 | 1.73484 | -0.379589 |
|  |  |  | Q6DHE2 | 1.73122 | -0.932569 |
|  |  |  | Q7ZVQ9 | 1.7306 | -0.53157 |
|  |  |  | Q8AX65 | 1.73049 | -1.24939 |
|  |  |  | Q6DGP6 | 1.72784 | -0.765021 |
|  |  |  | F1QZA5 | 1.7254 | -0.908111 |
|  |  |  | A0A2R8RYX8 | 1.72507 | 1.02307 |
|  |  |  | B3DGY6 | 1.72417 | 2.14314 |
|  |  |  | F1Q8A5 | 1.72287 | -0.817455 |
|  |  |  | F8W493 | 1.72235 | -0.763395 |
|  |  |  | E9QDU7 | 1.72185 | -0.786255 |
|  |  |  | Q7T3G2 | 1.71967 | -0.795235 |
|  |  |  | E9QI62 | 1.71854 | -0.528706 |
|  |  |  | E7FD28 | 1.71707 | -0.933865 |
|  |  |  | Q6P974 | 1.70983 | -3.62723 |
|  |  |  | Q50LC6 | 1.70904 | -1.34637 |
|  |  |  | Q7ZU04 | 1.70734 | -1.05722 |
|  |  |  | A3FKT8 | 1.70642 | -1.13953 |
|  |  |  | A0A0R4IJP7 | 1.69904 | -1.1636 |
|  |  |  | F1Q4Q8 | 1.69444 | -0.62413 |
|  |  |  | A0A2R8RLZ9 | 1.68942 | -0.824247 |
|  |  |  | Q9DGR6 | 1.68911 | -0.937433 |
|  |  |  | F1QML8 | 1.68849 | -0.613628 |
|  |  |  | Q7ZUY0 | 1.68849 | -0.486017 |
|  |  |  | F1RDQ9 | 1.68661 | -2.01399 |
|  |  |  | A8E5L0 | 1.68514 | -0.927063 |
|  |  |  | F1QNG7 | 1.68308 | -1.33081 |
|  |  |  | Q803T5 | 1.68266 | -0.908128 |
|  |  |  | Q7ZUA0 | 1.68265 | -0.9486 |
|  |  |  | Q9PWD8 | 1.68257 | -0.956165 |
|  |  |  | Q4V9I4 | 1.6819 | -0.680722 |
|  |  |  | A1L2C0 | 1.67993 | -2.00495 |
|  |  |  | A0A2R8QBA9 | 1.6782 | -0.604831 |
|  |  |  | Q9I9P6 | 1.67386 | -0.831944 |
|  |  |  | Q6DI13 | 1.67119 | -0.884586 |
|  |  |  | Q7ZV23 | 1.66799 | 1.14884 |
|  |  |  | Q6NXA6 | 1.66042 | -0.945162 |
|  |  |  | A7MCL3 | 1.65468 | -1.40826 |
|  |  |  | A0A2R8QIZ6 | 1.6529 | -1.07103 |
|  |  |  | Q7SXG5 | 1.65217 | -1.08307 |
|  |  |  | E7FCP7 | 1.65106 | -2.18885 |
|  |  |  | A0A0R4IF09 | 1.65046 | -0.617827 |
|  |  |  | A0A0R4IDD1 | 1.64825 | -0.758561 |
|  |  |  | B2GT73 | 1.64168 | -2.19592 |
|  |  |  | I3IRX8 | 1.63605 | -1.37299 |
|  |  |  | R4GEH4 | 1.63603 | -2.42831 |
|  |  |  | Q6DI07 | 1.63576 | -0.569078 |
|  |  |  | Q6DH18 | 1.63547 | -1.10329 |
|  |  |  | Q6P3J1 | 1.63526 | -0.673339 |
|  |  |  | Q803M8 | 1.63128 | -1.11561 |
|  |  |  | A2BHA3 | 1.63059 | -0.987551 |
|  |  |  | Q804W0 | 1.62952 | -1.51527 |
|  |  |  | A9JST0 | 1.62911 | -0.751495 |
|  |  |  | A8E7N8 | 1.62484 | -0.464805 |
|  |  |  | Q6PBX2 | 1.62405 | -0.965524 |
|  |  |  | B5DDE8 | 1.62375 | -0.292793 |
|  |  |  | E7FCN9 | 1.62337 | 1.04811 |
|  |  |  | A7MCA2 | 1.62263 | -0.307996 |
|  |  |  | A0A2R8RQC2 | 1.62164 | 0.867122 |
|  |  |  | Q801E6 | 1.62159 | -1.09706 |
|  |  |  | E7F8Y5 | 1.62135 | -0.856995 |
|  |  |  | Q802Z2 | 1.61566 | -0.726257 |
|  |  |  | B8A552 | 1.61543 | -0.614219 |
|  |  |  | Q6PC12 | 1.61439 | -0.966407 |
|  |  |  | Q7ZUI4 | 1.61438 | -0.960465 |
|  |  |  | F1RA36 | 1.61131 | -0.204753 |
|  |  |  | B2GT34 | 1.60861 | -0.382932 |
|  |  |  | C3S171 | 1.60767 | -0.782011 |
|  |  |  | Q6PHE9 | 1.60723 | -0.970661 |
|  |  |  | Q6IQN8 | 1.60595 | -0.820419 |
|  |  |  | B0S6C0 | 1.60312 | -1.28408 |
|  |  |  | B8JIS6 | 1.60111 | -1.78188 |
|  |  |  | Q7ZSY2 | 1.60061 | -1.41966 |
|  |  |  | F1R5I2 | 1.60034 | -0.976732 |
|  |  |  | Q504E8 | 1.59998 | -1.73613 |
|  |  |  | Q6NXA1 | 1.59593 | -0.519956 |
|  |  |  | Q5CZQ1 | 1.59138 | -1.35382 |
|  |  |  | A0A0R4IWE1 | 1.59042 | -1.34771 |
|  |  |  | A0A0R4IKF0 | 1.58877 | -0.705229 |
|  |  |  | A2CEF4 | 1.58717 | 3.20864 |
|  |  |  | Q6PFL9 | 1.58656 | -0.406681 |
|  |  |  | A9JRW0 | 1.58494 | -0.45687 |
|  |  |  | Q4VBI5 | 1.5843 | 0.476168 |
|  |  |  | A0A286YA42 | 1.58064 | -0.880753 |
|  |  |  | Q6PC35 | 1.57773 | -0.598238 |
|  |  |  | Q7SZD3 | 1.57747 | -0.349764 |
|  |  |  | X1WHF1 | 1.57559 | 0.397497 |
|  |  |  | A0A0R4II89 | 1.57353 | -0.402028 |
|  |  |  | Q3B7R1 | 1.5735 | -0.686892 |
|  |  |  | Q568G3 | 1.56915 | -0.863675 |
|  |  |  | A0A0R4IXJ1 | 1.56811 | -0.231111 |
|  |  |  | A0A2R8RK83 | 1.56224 | -1.11326 |
|  |  |  | B8JLJ3 | 1.56075 | -1.03291 |
|  |  |  | A5A4L9 | 1.55421 | 0.289016 |
|  |  |  | A0A0G2L4S8 | 1.548 | -1.1163 |
|  |  |  | F1R1E9 | 1.547 | -0.902438 |
|  |  |  | Q1L8Q3 | 1.54667 | 0.506811 |
|  |  |  | E7EZW7 | 1.54611 | -1.48681 |
|  |  |  | A0A0R4IPV5 | 1.53773 | -0.504957 |
|  |  |  | F1QZ40 | 1.53709 | -3.75906 |
|  |  |  | Q6YI49 | 1.53445 | -0.775828 |
|  |  |  | B0UYR8 | 1.53315 | -5.63875 |
|  |  |  | H0WE38 | 1.53275 | -1.095 |
|  |  |  | X1WDZ2 | 1.53046 | -3.0706 |
|  |  |  | Q6ZM23 | 1.52973 | 0.311012 |
|  |  |  | Q6DGX8 | 1.52774 | -0.516187 |
|  |  |  | A0A0R4IPH8 | 1.52736 | -1.32006 |
|  |  |  | Q804V8 | 1.52689 | -1.57198 |
|  |  |  | Q7ZVG1 | 1.52342 | -0.809268 |
|  |  |  | F8W4M7 | 1.52331 | -0.684727 |
|  |  |  | I3IS66 | 1.52304 | -0.998013 |
|  |  |  | E9QG01 | 1.5227 | 1.44444 |
|  |  |  | F1Q6S6 | 1.51804 | -1.88354 |
|  |  |  | F1R1S7 | 1.517 | 2.69122 |
|  |  |  | F1QY96 | 1.51687 | -0.41838 |
|  |  |  | Q7SXH8 | 1.51368 | -1.87264 |
|  |  |  | F1QJD1 | 1.51107 | -1.09308 |
|  |  |  | B2ZHD8 | 1.50993 | -2.98952 |
|  |  |  | E9QCE9 | 1.49533 | -0.847612 |
|  |  |  | A9JT49 | 1.49452 | -0.985494 |
|  |  |  | A7YYG7 | 1.49079 | -2.99221 |
|  |  |  | F1QEG3 | 1.48343 | -1.57734 |
|  |  |  | A0A2R8RUB5 | 1.47746 | -0.710625 |
|  |  |  | E9QJ96 | 1.47382 | -0.802926 |
|  |  |  | B2GRZ4 | 1.46709 | -0.90538 |
|  |  |  | Q5U7N6 | 1.46611 | -0.68398 |
|  |  |  | Q502M7 | 1.46532 | -1.98921 |
|  |  |  | Q7T2Q9 | 1.46465 | -0.449711 |
|  |  |  | A0A2R8Q860 | 1.46376 | -0.384536 |
|  |  |  | E9QGM5 | 1.45981 | -1.59585 |
|  |  |  | Q6GTE7 | 1.45962 | -0.705737 |
|  |  |  | Q52JI7 | 1.45741 | 1.07666 |
|  |  |  | F1R319 | 1.45151 | 0.792249 |
|  |  |  | A0A0R4IGG3 | 1.44962 | -1.02849 |
|  |  |  | Q7ZTT4 | 1.4464 | -1.19664 |
|  |  |  | Q5G9L7 | 1.44568 | -1.54851 |
|  |  |  | Q6P5L2 | 1.44435 | -1.09075 |
|  |  |  | A0A0R4IH39 | 1.44393 | -0.454965 |
|  |  |  | Q6PC38 | 1.44245 | -1.0796 |
|  |  |  | A8E585 | 1.44064 | -4.47252 |
|  |  |  | Q803J2 | 1.4359 | -0.963655 |
|  |  |  | Q5PQY4 | 1.43375 | -0.430407 |
|  |  |  | Q7T334 | 1.43366 | -0.626919 |
|  |  |  | F1QQQ5 | 1.43301 | -0.719625 |
|  |  |  | F1R314 | 1.4285 | 0.76864 |
|  |  |  | B8JIS8 | 1.42831 | -0.848928 |
|  |  |  | Q9DF44 | 1.42463 | -1.59278 |
|  |  |  | H1ZYM5 | 1.42453 | -1.1652 |
|  |  |  | Q7ZV76 | 1.4228 | 0.406517 |
|  |  |  | Q7T156 | 1.41923 | -1.88022 |
|  |  |  | A0A0R4IRN5 | 1.4189 | -1.15884 |
|  |  |  | Q6PBW4 | 1.41877 | -0.263955 |
|  |  |  | F1QP56 | 1.41559 | -0.898946 |
|  |  |  | A0A0R4I9C1 | 1.41516 | -1.0798 |
|  |  |  | F1RCA1 | 1.41324 | -0.763175 |
|  |  |  | F1R653 | 1.41322 | -0.370133 |
|  |  |  | Q6NYM4 | 1.41229 | -0.326156 |
|  |  |  | Q6TGV4 | 1.41162 | -0.262999 |
|  |  |  | A0A2R8QH08 | 1.41128 | -0.312284 |
|  |  |  | F6NTA0 | 1.41064 | -0.838587 |
|  |  |  | B3DJF3 | 1.41008 | -1.50957 |
|  |  |  | Q2YDP9 | 1.40995 | 0.987845 |
|  |  |  | B2GRW6 | 1.40957 | -0.958359 |
|  |  |  | Q5U3J6 | 1.40694 | 0.80883 |
|  |  |  | Q7T002 | 1.40404 | -0.532053 |
|  |  |  | Q7SYB4 | 1.39912 | -1.0933 |
|  |  |  | Q9PUL9 | 1.39816 | 0.269871 |
|  |  |  | Q6DGJ6 | 1.39693 | -0.597427 |
|  |  |  | F1QU55 | 1.39536 | -0.71823 |
|  |  |  | Q6IQR3 | 1.39398 | 3.12798 |
|  |  |  | Q7ZVR7 | 1.3937 | -1.11538 |
|  |  |  | A9JRA5 | 1.39202 | -1.05268 |
|  |  |  | Q6PBX7 | 1.39124 | -2.82339 |
|  |  |  | B3DJ97 | 1.3906 | -0.969916 |
|  |  |  | F1QCD4 | 1.38949 | -0.457088 |
|  |  |  | A0A0B5JW24 | 1.38838 | -1.31154 |
|  |  |  | F1R4Z8 | 1.38703 | -1.22689 |
|  |  |  | Q7SZP6 | 1.38619 | 1.7254 |
|  |  |  | Q803K3 | 1.38604 | -0.318909 |
|  |  |  | Q9PTF4 | 1.38445 | -0.836648 |
|  |  |  | A1L1W1 | 1.38435 | -0.501252 |
|  |  |  | F1QL66 | 1.38373 | -0.620274 |
|  |  |  | Q6PFS4 | 1.38036 | 0.567421 |
|  |  |  | Q5TZG5 | 1.37906 | -0.622723 |
|  |  |  | E7F5A4 | 1.3765 | -0.478291 |
|  |  |  | A0A0N4SUF2 | 1.37644 | -0.915457 |
|  |  |  | F1QUN8 | 1.37346 | 1.73253 |
|  |  |  | B5DDM9 | 1.37219 | -0.269396 |
|  |  |  | A0A0B5JPZ1 | 1.37112 | -0.894225 |
|  |  |  | A0A2R8QAS5 | 1.37013 | -0.412477 |
|  |  |  | Q7T3F3 | 1.36975 | -0.763591 |
|  |  |  | Q7ZUD3 | 1.36759 | -0.919408 |
|  |  |  | F1QFC0 | 1.36697 | -0.392207 |
|  |  |  | B0S6Z1 | 1.36692 | -0.42805 |
|  |  |  | A0A2R8QEY3 | 1.36642 | -0.838039 |
|  |  |  | B8A5D3 | 1.36615 | -0.889902 |
|  |  |  | A0A140LGQ3 | 1.36495 | -1.30568 |
|  |  |  | A7MC88 | 1.36442 | 2.28923 |
|  |  |  | F1QUS0 | 1.36406 | -1.55159 |
|  |  |  | A8WGS0 | 1.36373 | -0.863475 |
|  |  |  | Q4V8S1 | 1.36361 | -2.59733 |
|  |  |  | Q5RGA7 | 1.36294 | -1.49842 |
|  |  |  | Q6NYP6 | 1.36146 | -0.838071 |
|  |  |  | F8W4J1 | 1.35801 | -4.58485 |
|  |  |  | Q567X5 | 1.35759 | -2.55369 |
|  |  |  | F1R9Y5 | 1.35747 | -0.927973 |
|  |  |  | Q6NSN3 | 1.35728 | 0.325931 |
|  |  |  | F1QY43 | 1.35684 | -0.494035 |
|  |  |  | F1R5M2 | 1.35593 | -0.446496 |
|  |  |  | Q9I9E9 | 1.35451 | -0.592558 |
|  |  |  | A0A2R8QA71 | 1.35072 | -0.771836 |
|  |  |  | Q6P0I2 | 1.34824 | -0.585986 |
|  |  |  | A8BBA0 | 1.3479 | -0.881634 |
|  |  |  | F1R3I9 | 1.34597 | -0.627086 |
|  |  |  | Q4V8R9 | 1.34538 | -0.538153 |
|  |  |  | Q4KME8 | 1.34534 | -0.492283 |
|  |  |  | A0A0J9YJJ2 | 1.3363 | 0.841864 |
|  |  |  | B0S7W5 | 1.33548 | -1.00197 |
|  |  |  | Q5BJ14 | 1.33503 | 1.71655 |
|  |  |  | Q561U0 | 1.33319 | -0.748091 |
|  |  |  | Q6P3H9 | 1.33302 | 0.550823 |
|  |  |  | B2GRH9 | 1.33173 | -0.805693 |
|  |  |  | Q802D1 | 1.3293 | -0.501109 |
|  |  |  | Q801M0 | 1.32857 | -0.58626 |
|  |  |  | Q6PH48 | 1.32698 | -0.905899 |
|  |  |  | B3DKL0 | 1.32337 | -0.711011 |
|  |  |  | A4QN90 | 1.32087 | -0.164621 |
|  |  |  | A0A2R8QI78 | 1.32039 | -0.365772 |
|  |  |  | A4IG19 | 1.31719 | -0.715611 |
|  |  |  | Q5XIZ2 | 1.31125 | -1.17041 |
|  |  |  | Q6IQ94 | 1.311 | -1.82809 |
|  |  |  | F1Q6Z9 | 1.30886 | -0.710541 |
|  |  |  | Q2TV65 | 1.3065 | -0.526681 |
|  |  |  | Q5R2V3 | 1.30449 | 0.685958 |
|  |  |  | Q7SXW7 | 1.30168 | -0.865082 |
|  |  |  | F1R3F7 | 1.30012 | -0.32752 |
|  |  |  | A7Y224 | 1.29925 | 1.50364 |
|  |  |  | Q6TGU0 | 1.29734 | -0.594523 |
|  |  |  | B8A5M8 | 1.29664 | 1.51888 |
|  |  |  | B8JLQ3 | 1.29471 | -0.233745 |
|  |  |  | Q4VBK0 | 1.29453 | 0.478638 |

**Supplemental table 1**: Differentiated expressed proteins for the zebrafish larvae groups that were exposed to 2,4 DAB at 500µM up to 2 dpf (a) or up to 4 dpf (b).

| **Ingenuity Canonical Pathways** | **-log(p-value)** | **z-score** |
| --- | --- | --- |
| Protein Ubiquitination Pathway | 4.23 | NA |
| Pyrimidine Ribonucleotides De Novo Biosynthesis | 3 | -2.449 |
| Pyrimidine Ribonucleotides Interconversion | 2.87 | -2.236 |
| Aspartate Degradation II | 2.86 | -2 |
| Endoplasmic Reticulum Stress Pathway | 2.86 | -2 |
| HIF1α Signaling | 2.68 | -0.577 |
| Unfolded protein response | 2.5 | -2.646 |
| Role of PKR in Interferon Induction and Antiviral Response | 2.48 | 2.121 |
| ERK5 Signaling | 2.4 | -2.236 |
| Cell Cycle: G2/M DNA Damage Checkpoint Regulation | 2.4 | NA |
| BAG2 Signaling Pathway | 2.25 | -1.134 |
| Glucocorticoid Receptor Signaling | 2.13 | NA |
| FXR/RXR Activation | 2.02 | NA |
| IGF-1 Signaling | 2.02 | NA |
| MSP-RON Signaling In Cancer Cells Pathway | 2.02 | -1.89 |
| LXR/RXR Activation | 2.01 | -2.236 |
| Superpathway of Methionine Degradation | 2.01 | -2.449 |
| Hypoxia Signaling in the Cardiovascular System | 2.01 | NA |
| NRF2-mediated Oxidative Stress Response | 1.91 | -2.449 |
| Gluconeogenesis I | 1.88 | -2.828 |
| Aryl Hydrocarbon Receptor Signaling | 1.8 | 0 |
| Acute Phase Response Signaling | 1.8 | -2.236 |
| Aldosterone Signaling in Epithelial Cells | 1.8 | NA |
| eNOS Signaling | 1.79 | 2 |
| Amyotrophic Lateral Sclerosis Signaling | 1.77 | -0.447 |
| Maturity Onset Diabetes of Young (MODY) Signaling | 1.77 | NA |
| Xenobiotic Metabolism PXR Signaling Pathway | 1.71 | -2.111 |
| PPARα/RXRα Activation | 1.66 | 0.707 |
| Regulation of Cellular Mechanics by Calpain Protease | 1.66 | -2 |
| Xenobiotic Metabolism AHR Signaling Pathway | 1.66 | -1.134 |
| Glutathione Redox Reactions I | 1.61 | NA |
| Methionine Degradation I (to Homocysteine) | 1.61 | NA |
| Cysteine Biosynthesis III (mammalia) | 1.61 | NA |
| Vitamin-C Transport | 1.61 | NA |
| Apelin Adipocyte Signaling Pathway | 1.59 | -2.449 |
| Prostate Cancer Signaling | 1.55 | NA |
| NF-κB Signaling | 1.55 | -1.342 |
| Pyrimidine Deoxyribonucleotides De Novo Biosynthesis I | 1.53 | -2 |
| SPINK1 General Cancer Pathway | 1.53 | NA |
| Xenobiotic Metabolism Signaling | 1.49 | NA |
| Pyruvate Fermentation to Lactate | 1.43 | NA |
| Oleate Biosynthesis II (Animals) | 1.43 | NA |
| Thioredoxin Pathway | 1.43 | NA |
| D-glucuronate Degradation I | 1.43 | NA |
| GDP-L-fucose Biosynthesis I (from GDP-D-mannose) | 1.43 | NA |
| Rapoport-Luebering Glycolytic Shunt | 1.43 | NA |
| L-cysteine Degradation I | 1.43 | NA |
| Formaldehyde Oxidation II (Glutathione-dependent) | 1.43 | NA |
| Glutamate Degradation II | 1.43 | NA |
| Aspartate Biosynthesis | 1.43 | NA |
| Th17 Activation Pathway | 1.43 | NA |
| Glycolysis I | 1.36 | -2.646 |
| PI3K/AKT Signaling | 1.33 | -2.828 |
| RAN Signaling | 1.3 | -2 |
| Antioxidant Action of Vitamin C | 1.3 | NA |
| PPAR Signaling | 1.3 | 2 |

**Supplemental Table 2:** List of all enriched canonical pathway that were either activated or inhibited in the zebrafish larvae group that were exposed to 2,4 DAB at 500 µM until 4 days post fertilization (4 dpf) and their respective p values and z scores.

| **Gene Ontology (GO) Term** | **P Value** | **Fold Enrichment** |
| --- | --- | --- |
| **Metabolic processes** |  |  |
| GO:0009141~nucleoside triphosphate metabolic process | 0.003 | 7.946 |
| GO:0009123~nucleoside monophosphate metabolic process | 0.004 | 7.646 |
| GO:0042278~purine nucleoside metabolic process | 0.006 | 6.869 |
| GO:0009117~nucleotide metabolic process | 0.008 | 4.768 |
| GO:0006753~nucleoside phosphate metabolic process | 0.008 | 4.752 |
| GO:0009119~ribonucleoside metabolic process | 0.008 | 6.203 |
| GO:0009116~nucleoside metabolic process | 0.011 | 5.655 |
| GO:0055086~nucleobase-containing small molecule metabolic process | 0.014 | 4.121 |
| GO:0006163~purine nucleotide metabolic process | 0.015 | 5.195 |
| GO:0009259~ribonucleotide metabolic process | 0.016 | 5.087 |
| GO:0019693~ribose phosphate metabolic process | 0.017 | 4.942 |
| GO:0009132~nucleoside diphosphate metabolic process | 0.021 | 13.026 |
| GO:0019362~pyridine nucleotide metabolic process | 0.028 | 11.397 |
| GO:0006518~peptide metabolic process | 0.037 | 2.754 |
| GO:0006733~oxidoreduction coenzyme metabolic process | 0.040 | 9.233 |
| GO:0006002~fructose 6-phosphate metabolic process | 0.044 | 44.208 |
| GO:0006096~glycolytic process | 0.009 | 20.841 |
| **Developmental process** |  |  |
| GO:2000344~positive regulation of acrosome reaction | 0.001 | 56.110 |
| GO:0035036~sperm-egg recognition | 0.001 | 56.110 |
| GO:0007339~binding of sperm to zona pellucida | 0.001 | 56.110 |
| GO:0035803~egg coat formation | 0.001 | 56.110 |
| GO:0007340~acrosome reaction | 0.003 | 38.391 |
| GO:0007338~single fertilization | 0.006 | 24.315 |
| GO:0009566~fertilization | 0.008 | 22.104 |
| GO:0006096~glycolytic process | 0.009 | 20.841 |
| GO:0048477~oogenesis | 0.011 | 18.703 |
| GO:0007292~female gamete generation | 0.012 | 17.368 |
| GO:0007281~germ cell development | 0.026 | 11.765 |
| **Cellular** |  |  |
| GO:0006412~translation | 0.019 | 3.242 |
| GO:0043043~peptide biosynthetic process | 0.020 | 3.199 |
| GO:0043604~amide biosynthetic process | 0.030 | 2.914 |
| GO:0030036~actin cytoskeleton organization | 0.045 | 3.662 |
| GO:0034622~cellular macromolecular complex assembly | 0.053 | 2.872 |
| **Other** |  |  |
| GO:0044724~single-organism carbohydrate catabolic process | 0.024 | 12.157 |

**Supplemental table 3:** List enriched of GO Biological Processes derived from the differentiated expressed proteins in the zebrafish larvae groups that were exposed to 2,4 Dab at 500 µM until 2 days post fertilization (2 dpf).

| **Gene Ontology (GO) Term** | **PValue** | **Fold Enrichment** |
| --- | --- | --- |
| **Metabolic processes** |  |  |
| GO:0009132~nucleoside diphosphate metabolic process | 0.0000 | 11.9402 |
| GO:0019752~carboxylic acid metabolic process | 0.0000 | 3.7147 |
| GO:0043436~oxoacid metabolic process | 0.0000 | 3.6725 |
| GO:0009116~nucleoside metabolic process | 0.0000 | 4.6650 |
| GO:0009119~ribonucleoside metabolic process | 0.0000 | 4.8329 |
| GO:0042278~purine nucleoside metabolic process | 0.0000 | 5.0369 |
| GO:0055086~nucleobase-containing small molecule metabolic process | 0.0000 | 3.4629 |
| GO:0009141~nucleoside triphosphate metabolic process | 0.0000 | 5.0986 |
| GO:0009117~nucleotide metabolic process | 0.0000 | 3.4598 |
| GO:0006753~nucleoside phosphate metabolic process | 0.0000 | 3.4485 |
| GO:0006163~purine nucleotide metabolic process | 0.0000 | 3.8100 |
| GO:0009259~ribonucleotide metabolic process | 0.0000 | 3.7303 |
| GO:0019693~ribose phosphate metabolic process | 0.0000 | 3.6241 |
| GO:0009123~nucleoside monophosphate metabolic process | 0.0001 | 4.2053 |
| GO:0019362~pyridine nucleotide metabolic process | 0.0001 | 6.9651 |
| GO:0006733~oxidoreduction coenzyme metabolic process | 0.0005 | 5.6426 |
| GO:0006220~pyrimidine nucleotide metabolic process | 0.0019 | 9.2868 |
| GO:0019318~hexose metabolic process | 0.0040 | 5.6665 |
| GO:0005996~monosaccharide metabolic process | 0.0060 | 5.1435 |
| GO:0006213~pyrimidine nucleoside metabolic process | 0.0146 | 7.6856 |
| GO:0046500~S-adenosylmethionine metabolic process | 0.0157 | 15.1966 |
| GO:0009112~nucleobase metabolic process | 0.0280 | 6.0239 |
| GO:0006631~fatty acid metabolic process | 0.0312 | 2.9549 |
| GO:0006096~glycolytic process | 0.0000 | 11.1442 |
| **Developmental Processes** |  |  |
| GO:0001654~eye development | 0.0250 | 2.0520 |
| GO:0043010~camera-type eye development | 0.0320 | 2.1506 |
| GO:0034728~nucleosome organization | 0.0146 | 3.5139 |
| GO:0060218~hematopoietic stem cell differentiation | 0.0224 | 6.5554 |
| GO:0006334~nucleosome assembly | 0.0335 | 3.3433 |
| GO:0031497~chromatin assembly | 0.0400 | 3.1840 |
| **Cellular Processes** |  |  |
| GO:0009163~nucleoside biosynthetic process | 0.0015 | 4.7422 |
| GO:1901659~glycosyl compound biosynthetic process | 0.0016 | 4.6434 |
| GO:0072528~pyrimidine-containing compound biosynthetic process | 0.0027 | 8.4426 |
| GO:0090407~organophosphate biosynthetic process | 0.0029 | 2.7335 |
| GO:0044744~protein targeting to nucleus | 0.0040 | 5.6665 |
| GO:1902593~single-organism nuclear import | 0.0040 | 5.6665 |
| GO:0006606~protein import into nucleus | 0.0040 | 5.6665 |
| GO:0072522~purine-containing compound biosynthetic process | 0.0045 | 3.4585 |
| GO:0009165~nucleotide biosynthetic process | 0.0047 | 3.1304 |
| GO:1901293~nucleoside phosphate biosynthetic process | 0.0053 | 3.0785 |
| GO:0017038~protein import | 0.0058 | 4.2862 |
| GO:0072594~establishment of protein localization to organelle | 0.0115 | 3.2538 |
| GO:0006913~nucleocytoplasmic transport | 0.0124 | 3.6453 |
| GO:0051169~nuclear transport | 0.0124 | 3.6453 |
| GO:0006333~chromatin assembly or disassembly | 0.0135 | 3.5784 |
| GO:0046390~ribose phosphate biosynthetic process | 0.0138 | 3.1392 |
| GO:0045333~cellular respiration | 0.0180 | 3.9332 |
| GO:0009142~nucleoside triphosphate biosynthetic process | 0.0210 | 4.7221 |
| GO:0006605~protein targeting | 0.0306 | 2.6693 |
| GO:0046112~nucleobase biosynthetic process | 0.0324 | 10.4477 |
| GO:1902582~single-organism intracellular transport | 0.0332 | 2.6222 |
| GO:0071103~DNA conformation change | 0.0353 | 2.8680 |
| GO:0046166~glyceraldehyde-3-phosphate biosynthetic process | 0.0354 | 55.7208 |
| GO:0065004~protein-DNA complex assembly | 0.0364 | 2.8470 |
| GO:0033365~protein localization to organelle | 0.0390 | 2.5328 |
| GO:0009226~nucleotide-sugar biosynthetic process | 0.0445 | 8.7980 |
| GO:0006869~lipid transport | 0.0446 | 2.7087 |
| **Other** |  |  |
| GO:0044724~single-organism carbohydrate catabolic process | 0.0000 | 8.3581 |
| GO:0030163~protein catabolic process | 0.0002 | 2.7049 |
| GO:0044275~cellular carbohydrate catabolic process | 0.0002 | 16.3885 |
| GO:0051603~proteolysis involved in cellular protein catabolic process | 0.0002 | 2.7008 |
| GO:0044257~cellular protein catabolic process | 0.0003 | 2.6802 |
| GO:0043632~modification-dependent macromolecule catabolic process | 0.0012 | 2.6299 |
| GO:0044265~cellular macromolecule catabolic process | 0.0028 | 2.1829 |

**Supplemental table 4:** List enriched of GO Biological Processes derived from the differentiated expressed proteins in the zebrafish larvae groups that were exposed to 2,4 Dab at 500 µM until 4 days post fertilization (4 dpf).

| **Symbol** | **Entrez Gene Name** | **UniProt/Swiss-Prot Accession** | **Expr p-value** | **Expr Log Ratio** |
| --- | --- | --- | --- | --- |
| ANP32B | acidic nuclear phosphoprotein 32 family member B | F1Q4N7 | 0.008 | -0.992 |
| COPE | COPI coat complex subunit epsilon | F1R7S1 | 0.010 | -0.535 |
| DLST | dihydrolipoamide S-succinyltransferase | Q6NZW7 | 0.033 | 1.402 |
| DYNLL2 | dynein light chain LC8-type 2 | Q7SZP6 | 0.044 | 1.986 |
| EEF1G | eukaryotic translation elongation factor 1 gamma | Q8JIU6 | 0.016 | -0.498 |
| EEF2 | eukaryotic translation elongation factor 2 | Q6P3J5 | 0.039 | -0.306 |
| EFTUD2 | elongation factor Tu GTP binding domain containing 2 | F1Q6N0 | 0.046 | -0.390 |
| EWSR1 | EWS RNA binding protein 1 | Q6NUX1 | 0.020 | -0.411 |
| GANAB | glucosidase II alpha subunit | F1Q6Z9 | 0.020 | -0.914 |
| GYG1 | glycogenin 1 | Q803Q1 | 0.046 | -0.809 |
| HP1BP3 | heterochromatin protein 1 binding protein 3 | A0A140LGU1 | 0.005 | 1.365 |
| KTN1 | kinectin 1 | F1QJA1 | 0.007 | 2.847 |
| LAMP1 | lysosomal associated membrane protein 1 | E9QCP4 | 0.009 | -1.290 |
| LIN7C | lin-7 homolog C, crumbs cell polarity complex component | A0A2R8QIA2 | 0.008 | 1.769 |
| LMNB2 | lamin B2 | B3DFN3 | 0.033 | 0.264 |
| MDH2 | malate dehydrogenase 2 | Q7T334 | 0.028 | -0.722 |
| OSTC | oligosaccharyltransferase complex non-catalytic subunit | A0A0R4I9P9 | 0.046 | -0.492 |
| PDIA4 | protein disulfide isomerase family A member 4 | Q6P3I1 | 0.032 | -0.856 |
| PLEC | plectin | A0A2R8QLG6 | 0.047 | 1.963 |
| PPIA | peptidylprolyl isomerase A | B8JKN6 | 0.014 | -1.158 |
| PPIB | peptidylprolyl isomerase B | Q6PBW4 | 0.018 | -0.457 |
| PRDX4 | peroxiredoxin 4 | F1QSP8 | 0.001 | -1.228 |
| PRKCSH | protein kinase C substrate 80K-H | Q802Z2 | 0.029 | -0.772 |
| PTGR1 | prostaglandin reductase 1 | A0A2R8RPQ9 | 0.007 | -1.590 |
| PYGB | glycogen phosphorylase B | A4IG19 | 0.023 | -0.931 |
| RHOA | ras homolog family member A | A7MCP4 | 0.042 | 0.286 |
| RPN1 | ribophorin I | Q8AYB9 | 0.012 | -1.458 |
| RPN2 | ribophorin II | Q7ZV07 | 0.012 | -0.346 |
| SEC13 | SEC13 homolog, nuclear pore and COPII coat complex component | A0A2R8RKG2 | 0.034 | -0.491 |
| SLC25A11 | solute carrier family 25 member 11 | F1R319 | 0.012 | 1.112 |
| SMYD1 | SET and MYND domain containing 1 | Q2MJQ9 | 0.014 | 1.021 |
| SNX6 | sorting nexin 6 | F8W5C6 | 0.003 | 2.240 |
| **SOD1** | **superoxide dismutase 1** | **B2GRH9** | **0.012** | **-1.413** |
| TKT | transketolase | E7F8S4 | 0.007 | 3.172 |
| VAT1 | vesicle amine transport 1 | A7YYF7 | 0.000 | 3.804 |

**Supplemental table 5:** Differentially expressed proteins found to be up/downregulated in 2,4 DAB exposed larval zebrafish compared to control (Top score 68).

|  | **Minus Log of P Value** | | **Z Score** | |
| --- | --- | --- | --- | --- |
| **Canonical Pathways (CTRL vs 2,4 DAB)** | **CTRL** | **2,4 DAB** | **CTRL** | **2,4 DAB** |
| EIF2 Signaling | 2.0 | 9.9 | -4.4 | -5.0 |
| Coronavirus Pathogenesis Pathway | 1.1 | 7.1 | 3.5 | 4.7 |
| Glycolysis I | 3.0 | 2.5 | 3.7 | 1.5 |
| Gluconeogenesis I | 2.3 | 2.5 | 3.6 | 1.5 |
| Calcium Signaling | 1.5 | 2.2 | 2.3 | 2.6 |
| Regulation of eIF4 and p70S6K Signaling | 0.0 | 3.5 | 2.1 | 0.6 |
| mTOR Signaling | 0.0 | 3.1 | 2.6 | N/A |
| Phototransduction Pathway | 2.5 | 0.6 | N/A | 1.0 |
| Sucrose Degradation V (Mammalian) | 0.8 | 2.2 | N/A | N/A |
| Glutamate Receptor Signaling | 0.3 | 2.2 | N/A | N/A |
| 14-3-3-mediated Signaling | 2.2 | 0.0 | N/A | N/A |
| Cellular Effects of Sildenafil (Viagra) | 1.5 | 0.3 | N/A | N/A |
| Protein Kinase A Signaling | 1.3 | 0.5 | N/A | N/A |
| TR/RXR Activation | 0.0 | 1.6 | N/A | N/A |
| Protein Ubiquitination Pathway | 0.0 | 1.4 | N/A | N/A |

**Supplemental table 6:** All categories of enriched canonical in the systemic analysis for the control and 2,4 DAB treated zebrafish larvae along with their associated minus log of p value and z-scores.

| **Upstream Regulators**  **(CTRL vs 2,4 DAB)** | **CTRL** | **2,4 DAB** |
| --- | --- | --- |
| LARP1 | 5.606 | 6.63 |
| MLXIPL | -5.191 | -6.564 |
| MYCN | -3.978 | -5.409 |
| MYC | -2.259 | -5.868 |
| SMARCA4 | 3.283 | 2.84 |
| MEF2C | 2.541 | 3.067 |
| beta-estradiol | 2.536 | -3.068 |
| sirolimus | 0.926 | 4.613 |
| RICTOR | N/A | 5.175 |
| BDNF | 2.334 | 2.796 |
| ISL1 | -2.646 | -2.236 |
| NOS2 | -2.201 | -2.333 |
| ARNT2 | 2.496 | 1.89 |
| DNMT3B | -2.111 | -2.121 |
| SIM1 | 2.309 | 1.89 |
| TBX5 | 1.586 | 2.592 |
| 5-fluorouracil | N/A | 4.126 |
| PHA-666859 | -2 | -2 |
| DMD | 2.333 | 1.633 |
| 1,2-dithiol-3-thione | N/A | -3.881 |
| torin1 | -3.178 | -0.687 |
| valproic acid | 1.793 | 2.039 |
| ADAM12 | -2.433 | -1.387 |
| SRF | 1.927 | 1.731 |
| NFE2L2 | N/A | -3.52 |
| tanespimycin | 1.5 | 1.994 |
| phenylbutazone | 1.499 | -1.98 |
| FMR1 | -1.885 | -1.29 |
| poly rI:rC-RNA | N/A | -3.094 |
| EPO | -1.4 | -1.664 |
| pirinixic acid | N/A | -3.041 |
| bortezomib | N/A | -3.04 |
| TCR | N/A | -2.86 |
| UBQLN2 | N/A | 2.714 |
| RTN4 | -1 | 1.633 |
| lipopolysaccharide | 0.071 | -2.526 |
| CTNNB1 | 1.401 | 1.195 |
| TFAP2A | N/A | 2.566 |
| N-nitro-L-arginine methyl ester | -2.19 | -0.277 |
| BCR-ABL1 | 2.449 | N/A |
| FAAH | N/A | 2.449 |
| LDL | N/A | -2.432 |
| 2,3 butanedione monoxime | N/A | 2.415 |
| progesterone | 2.397 | N/A |
| prostaglandin J2 | N/A | -2.376 |
| mir-210 | -2.343 | N/A |
| MXD1 | N/A | 2.335 |
| uranyl nitrate | -1.171 | -1.127 |
| D-glucose | 2.056 | 0.211 |
| Gm15807/Hmgn5 | N/A | 2.236 |
| SOX7 | -2.236 | N/A |
| MUC1 | N/A | -2.226 |
| GATA4 | 0.969 | 1.248 |
| cobalt chloride | 1.237 | 0.975 |
| CRX | 2.2 | N/A |
| PSEN1 | -1.264 | -0.933 |
| SRC | 2.178 | N/A |
| dihydrotestosterone | 2.106 | 0.06 |
| SREBF1 | 0.608 | -1.548 |
| NFE2L1 | N/A | -2.138 |
| SMTNL1 | N/A | -2.121 |
| ciprofloxacin | -2.065 | N/A |
| KDM8 | 2.01 | N/A |
| BCL2 | 1.219 | 0.784 |
| SNCA | N/A | -2 |
| MED13 | -2 | N/A |
| EPHB4 | 1 | 1 |
| NGFR | 1 | 1 |
| enalapril | N/A | -2 |
| MAF | N/A | 2 |
| TP53RK | N/A | -1.982 |
| BO-653 | N/A | 1.982 |
| probucol | N/A | 1.982 |
| deoxycholate | N/A | -1.981 |
| potassium chloride | N/A | 1.98 |
| APOA1 | N/A | -1.969 |
| VIP | 1.964 | N/A |
| PPARG | 1.946 | N/A |
| MYOD1 | N/A | 1.915 |
| KAT5 | N/A | -1.912 |

**Supplemental table 7**: All categories of upstream regulators

predicted to be activated or inhibited in the systemic analysis.

for the control and 2,4 DAB treated zebrafish larvae along with

their associated z-scores.
